# Dysfunctional memory B cell responses to the protective repeat region of the *Plasmodium* circumsporozoite protein are associated with waning humoral immunity

**DOI:** 10.64898/2025.12.19.695559

**Authors:** Courtney E McDougal, Catherine G Mkindi, Lauren B Rodda, Caroline V Lucarelli, Mark D Langowski, Neil P King, Said Jongo, Claudia Daubenberger, Marion Pepper

## Abstract

Malaria vaccines provide waning protection from disease that is correlated with the production of antibodies to the repeat region of the circumsporozoite protein (CSP). CSP-based vaccines display limited durability in malaria-naïve individuals yet are even less effective in malaria-experienced individuals, suggesting the generation of non-optimal humoral immunity in response to both infection and vaccination. To address this hypothesis, we performed a cross-species, comprehensive analysis of B cell responses to CSP after *Plasmodium* infection or immunization, focusing our analysis on the repeat and C-terminus domains included in malaria subunit vaccines. Herein we demonstrate that the repetitive nature of the protective region of the CSP protein independently impacts the differentiation of the CSP-specific B cells, impinging on their ability to recall for multiple subsequent exposures.

## Introduction

Over 50 years of research and significant investment resulted in the first WHO-recommended malaria vaccine, RTS,S/AS01^1^, targeting the *Plasmodium* circumsporozoite protein (CSP). RTS,S has revolutionized immune-mediated control of malaria infection, leading to upwards of 80% protection in controlled human malaria infection trials and a reported 13% reduction in mortality since widespread use in late 2021^2^, potentially saving tens of thousands of lives annually. However, despite the early success, a significant challenge remains for CSP-based vaccines: the rapid waning of immune protection^3–5^. Efficacy of RTS,S drops dramatically by 18 months after immunization and is associated with a steep decline in anti-CSP antibody titers. Boosters can restore some efficacy, though the level of anti-CSP serum antibody never reaches titers detected after initial immunization and, eventually, yearly immunizations no longer boost antibody titers at all^4,6^. This leads to efficacy plummeting to a mere 4% after seven years and a negative correlation with protection over time^3^. This is in stark contrast to other vaccines, such as yellow fever^7^ or smallpox^8^, where protection can last decades. An additional challenge of CSP-based vaccines is a loss of efficacy in individuals living in malaria-endemic areas, again largely associated with diminished antibody titers^4,5^. In one study, median CSP-specific antibody responses generated by vaccination were almost five times lower in Tanzanian vaccine recipients compared to malaria-naïve American vaccine recipients^9^. Areas of high transmission, such as Bamako, Mali, demonstrate even more impaired antibody responses when compared to Tanzanian participants in areas of lower endemicity^9^. Although it is clear that humoral immunity and CSP-based vaccine efficacy wanes especially in areas of highest endemicity, it is not clear which factors impinge upon the generation of long-lasting CSP-specific humoral immunity.

To reveal the underlying mechanisms associated with a loss of humoral immunity to CSP-based vaccines, we performed an in-depth analysis of the development of CSP-specific B cell responses in both mice and humans. We focused on two domains of CSP that are included in the RTS,S vaccine, the immunodominant repeat region (RR) and the non-repetitive C-terminus domain (CT)^10–12^. The CT contains T cell epitopes that may be important for T-dependent help^13–15^ despite CT-specific B cell responses being infrequently associated with protection^16,17^. In contrast, induction of RR-specific antibody is critical for vaccine efficacy and correlates with protection across multiple studies and vaccine modalities^12,18,19^. While such data suggest that immune responses to the RR can be protective, other data suggest that the parasite uses the RR to evade productive immune responses^10,20^. It has been proposed that the repetitive nature of the RR may function as an immunodominant decoy, preventing immune response towards other, potentially more protective epitopes^10,20^. Consistent with this hypothesis, immunization with repeat-truncated versions of CSP generates higher antibody titers towards subdominant epitopes and leads to higher levels of protection from challenge^10^.

In addition, previous work has demonstrated that the repetitive nature of the RR allows for B cell receptor (BCR) crosslinking^21^. BCR crosslinking influences intracellular signaling pathways and shapes B cell differentiation choices^22^. While some BCR crosslinking can be beneficial by enhancing early antibody-secreting plasmablasts (PBs)^23^, excessive signaling can also negatively impact the formation of durable memory populations. For example, B cell binding to a repetitive 23-valent pneumococcal polysaccharide vaccine leads to aberrant BCR crosslinking and a subsequent failure to generate long-lived memory populations^24^. We therefore hypothesize that the repetitive nature of the RR impacts B cell differentiation and, ultimately, the durability of RR-specific memory populations.

To test this hypothesis, we generated B cell tetramers containing either the RR or the CT of the CSP protein to characterize the B cell pool established after infection and/or immunization in both mouse and human samples. We show that *Plasmodium* infection generates a population of early RR-specific short-lived PBs and later memory B cells (MBCs) that are phenotypically-associated with a functional population of MBCs. Upon rechallenge with a murine model of RTS,S, these MBCs differentiate optimally into antibody-secreting PBs. However, further analysis reveals that differentiation into these short-lived effectors upon recall is at the expense of replenishing the long-lived memory pool. Failure to replenish RR-specific memory pools leads to a depletion of RR-specific B cells that is exacerbated with additional exposures to the CSP antigen. Using novel atomically-designed protein-based repeat-truncated nanoparticles, we demonstrate that RR valency and antigen density specifically and independently contribute to impaired MBC differentiation. Human PBMCs from malaria-experienced adults similarly lacked functional RR-specific memory and, in contrast to *Plasmodium*-naïve individuals, did not boost RR-specific responses upon vaccination. Interestingly, vaccination of children in endemic areas, who have likely been infected fewer times than adults, led to superior B cell responses that mimic vaccination of malaria-naïve adults. Overall, our data in human samples and murine models reveal that the repetitive nature of the RR influences B cell differentiation. Specifically, we show for the first time that while RR-specific MBCs can differentiate into functional, short-lived effectors upon recall, they fail to renew durable memory pools. This provides temporary protection but ultimately leads to impaired long-lived immunity.

## Results

### Following *Plasmodium* infection, RR-specific - but not CT-specific - B cells differentiate into germinal center-independent memory B cells

As CSP-vaccine induced immune protection wanes in response to both *Plasmodium* infection and CSP-based vaccination, we hypothesized that perhaps this was due to an aberrant B cell response driven by the CSP antigen itself, and more specifically the protective repeat region. We therefore first assessed the kinetics and phenotypic attributes of B cells specific for the RR or the CT of the CSP protein after wild-type *Plasmodium* infection. C57BL/6 mice were infected with 2000 *Plasmodium yoelii* sporozoites that express the *Plasmodium falciparum* CSP (*Py^Pf^*^CSP^) instead of the endogenous CSP protein^25^, as this allowed us to use the same reagents to assess CSP-specific responses in both a murine model and, later, in human clinical samples. Domain-specific B cells were isolated from spleens at early, peak, and memory time points (seven, fourteen, and at least sixty days later) (Fig. 1a and Extended Data Fig. 1). While RR-specific cells expanded and differentiated into short-lived, antibody-producing CD138^+^IgM^+^ PBs (Fig. 1b-c, Extended Data Fig. 2a-c) and a CD38^+^GL7^+^ high germinal center (GC)-precursor population (Fig. 1d), a similarly robust response was not observed to the CT (Fig. 1b-d). Furthermore, the RR-specific GC-precursor population did not fully commit to a CD38^-^GL7^+^ GC B cell fate at any timepoint examined (Fig. 1d), consistent with previous work from our lab examining the total CSP-specific response to wild-type *Plasmodium* infection^26^.

**Fig. 1:**
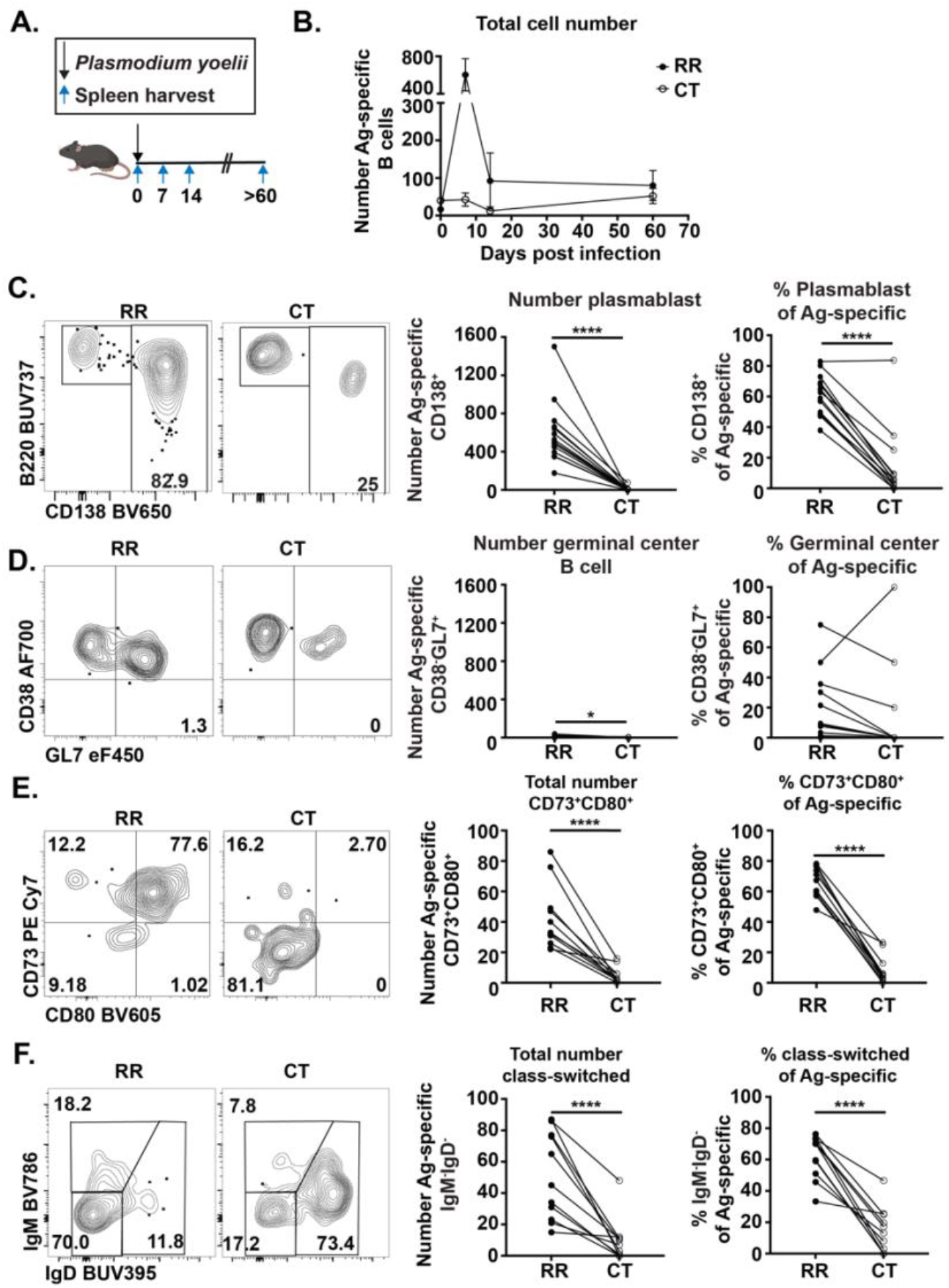
Following *Plasmodium* infection, RR-specific B cells—but not CT-specific B cells—differentiate into germinal center-independent memory B cells. **a,** C57BL/6 mice were infected with 2000 *Py^Pf^*^CSP^ sporozoites. Seven, fourteen, and at least sixty days later, splenic RR- and CT-specific B cells were isolated and characterized. **b,** The total number of CT- and RR-specific B cells throughout the time course. **c,** RR- and CT-specific PBs (CD138^+^) 7 days post infection. **d,** RR-and CT-specific GC-precursors (CD38^+^GL7^+^) and GC B cells (CD38^-^GL7^+^) 14 days post infection. **e,** Expression of CD73 and CD80 on RR- or CT- specific B cells at day 60 post-infection. **f,** Class-switched (IgD^-^IgM^-^) RR- or CT-specific B cells at day 60 post-infection. Gates were drawn based on naïve controls. n = 11-13 mice per timepoint. Statistical analysis by paired t test *p<0.05 ****p<0.0001

It was unclear, however, if the CD38^+^GL7^+^ RR-specific GC precursor population we observed in Fig. 1d could eventually form a long-lived memory population in response to infection as there was no indication of GC formation. We therefore assessed the phenotype of the RR- and CT- specific MBCs, focusing on BCR isotype and the expression of surface proteins associated with specific functional attributes of MBCs during a recall response including CD80 and CD73^27–31^. Specifically, IgG^+^ or IgM^+^ CD73^+^ CD80^+^ MBCs preferentially form short-lived, antibody-secreting PBs upon antigen restimulation that are important for immediate protection^27–31^. In contrast, IgM^+^ or IgD^+^ MBCs that express one or fewer of these markers preferentially form secondary GCs^27–31^. Surprisingly, after at least sixty days post-infection, we could find a population of RR-specific MBCs, most of which were class-switched and expressed both CD73 and CD80 (Figs. 1e,f). In contrast, most CT-specific B cells appeared naïve, as few were class-switched and most did not express CD73 or CD80 (Figs. 1e,f). Taken together, these data demonstrate that wild-type *Plasmodium* infection generates RR-specific MBCs, but not CT-specific MBCs. Furthermore, despite the lack of GCs during wild-type infection, RR-specific B cells form a class-switched population with high expression of CD73 and CD80 that predicts strong functional responses to a subsequent challenge.

### Following CSP-vaccine immunization, RR- and CT-specific B cell responses differentiate in distinct ways and only RR-specific MBCs can form in a GC- independent manner

To determine if the inability of RR-specific B cells to enter GCs was an infection-specific phenomenon or a common attribute of responses to that specific region of CSP, we assessed RR-specific GC responses in two immunization models. First, we immunized mice with 25*μ*g of an icosahedral two-component nanoparticle, RT-I53-50^32–34^, adjuvanted with Sigma Adjuvant System (Fig. 2a). RT-I53-50 displays 60 copies of the antigenic portion of the RTS,S malaria vaccine, which consists of 19 repeats of the RR amino acid sequence “NANP” and the entire CT domain^26^. Second, we immunized mice with irradiated *Py^Pf^*^CSP^ (Extended Data Fig. 3a). Irradiation prevents progression to blood-stage and generates robust GCs to CSP broadly^26^, though whether these are specific to the RR domain is unclear. In both immunization models, many CSP RR-specific PBs were detected seven days post-immunization with a small GC B population forming one week later (Fig. 2b-c, Extended Data Fig. 3b-c). Further analysis revealed that immunization with RT-I53-50 or irradiated *Py^Pf^*^CSP^ also induced a robust expansion of CT-specific PBs and GCs, demonstrating that the lack of B cell expansion to the CT we see in infection is not due to an inability of our reagents to identify sporozoite-expressed CSP CT (Figs. 2a-c, Extended Data Figs. 3b-c). Importantly, though both CT- and RR-specific responses were generated in these immunization models, RR-specific B cells were significantly more likely to differentiate into PBs, whereas CT-specific B cells preferentially entered GCs (Figs. 2b-c, Extended Data Figs. 3b-c). Taken together with our above data from the infection model, these data demonstrate that a) even in an immunization model, distinct responses form to distinct regions of the same protein suggesting that repetitive epitopes may polarize B cells differently than monomeric epitopes; b) although infection has a more profoundly negative impact on RR-and CT-specific GC formation than immunization, CD80^+^CD73^+^ cells RR-specific can form in response to either model; and c) raises the formal possibility that CD80^+^CD73^+^ CSP RR-specific MBCs can form independently of a GC in response to either infection or immunization.

**Fig. 2:**
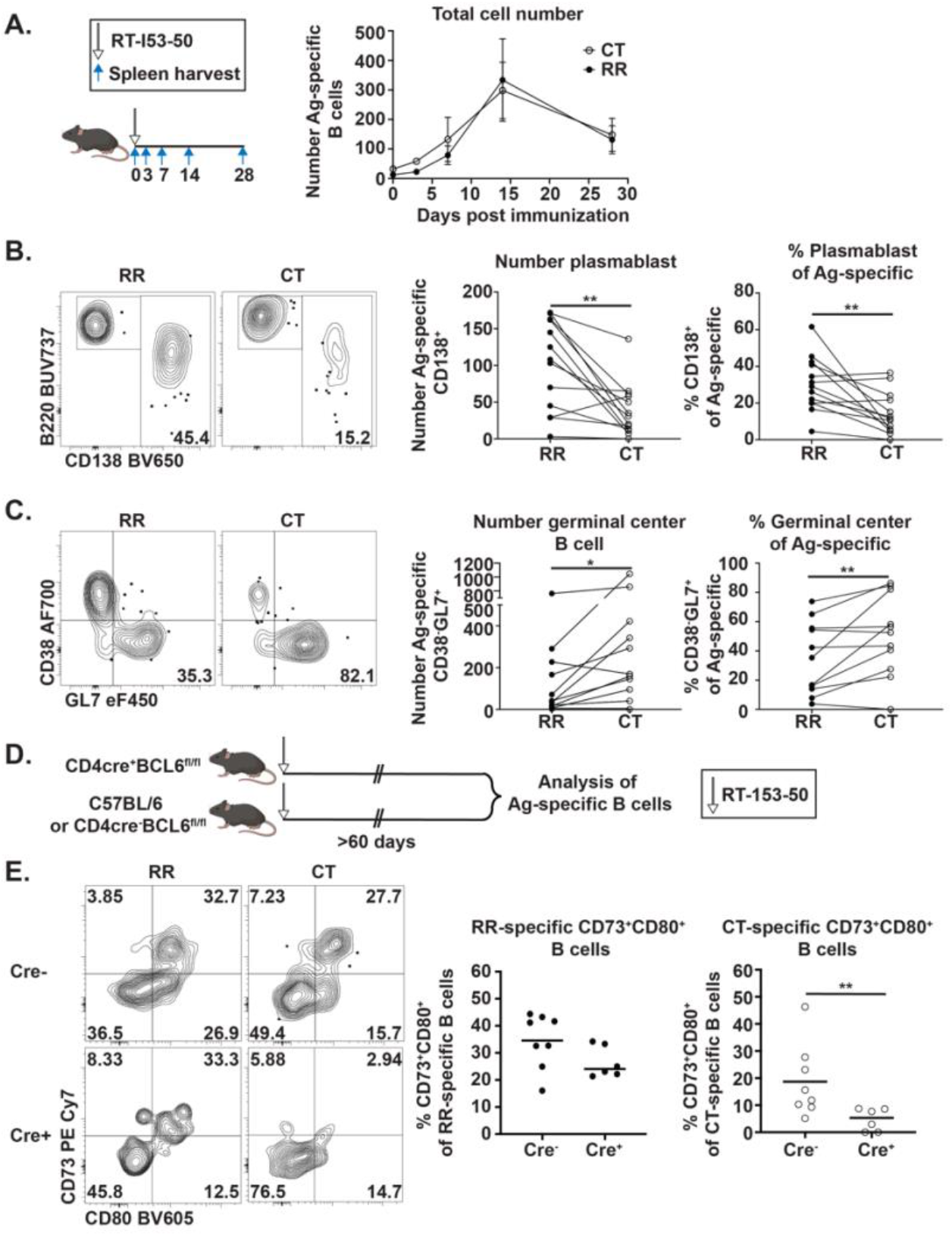
After CSP immunization, CT-specific B cells preferentially differentiate into GCs, whereas RR-specific B cells favor polarization toward short-lived PBs and MBCs that do not require GCs. **a,** C57BL/6 mice were immunized with 25*µ*g RT-I53-50 intraperitoneally. Spleens were harvested at indicated days post immunization. The total number of RR- and CT-specific B cells at indicated times after immunization are shown. **b,** RR- and CT-specific PBs (CD138^+^) at day 7 post RT-I53-50 immunization. **c,** RR- and CT-specific GCs (CD38^-^GL7^+^) at day 14 post RT-I53-50 immunization. **d,** Bcl6^fl/fl^ CD4cre^+^ or control mice (Bcl6^fl/fl^ CD4cre^-^ and C57BL/6) were immunized with 25*µ*g RT-I53-50 intraperitoneally. Splenic CT- and RR-specific B cells were isolated and characterized at least 60 days later. **e,** CD73 and CD80 expression on RR- and CT-specific B cells at day 60 post immunization in indicated mouse strains. Gates were drawn based on naïve controls. Statistical analysis by paired t test (b,c) or Mann-Whitney (e). n = 8-13 mice. *p<0.05 **p<0.01

To test this possibility, we immunized CD4cre^+^ Bcl6^fl/fl^ mice, which lack GC T follicular helper cells (and therefore GCs), allowing us to assess formation of GC-independent B cell memory. We immunized CD4cre^+^ Bcl6^fl/fl^ experimental mice and CD4cre^-^ Bcl6^fl/fl^ or C57BL/6 wild-type control mice with 25*μ*g RT-I53-50 and examined B cell differentiation in each scenario (Fig. 2d). In control mice, we detected a population of both CT- and RR-specific CD73^+^CD80^+^ MBCs 60 days after RT-I53-50 immunization (Fig. 2e). In contrast, in CD4cre^+^ Bcl6^fl/fl^ mice, we observed comparable numbers of RR-specific CD73^+^CD80^+^ MBCs to control mice, whereas there was a significant loss of CT-specific CD73^+^CD80^+^ MBCs compared to the WT setting (Fig. 2e). This demonstrates that the RR antigen is capable of generating CD73^+^CD80^+^ MBCs even in the absence of a GC, whereas the non-repetitive CT antigen requires a GC for the generation of CD73^+^CD80^+^ MBCs.

As GCs were not critical for the expansion of RR-specific B cells, we next asked whether T cells broadly are required. RR and CT-specific B cell responses in C57BL/6 or TCRα^-/-^ mice immunized with 25*μ*g RT-I53-50 were assessed seven days later (Extended Data Fig. 4a). Expansion of both CT- and RR-specific B cells was impaired in the absence of T cells compared to control mice, as was differentiation into PBs (Extended Data Figs. 4b,c). This is consistent with previous research demonstrating that RR-specific responses generated after an immunization with irradiated *Plasmodium berghei* require T cells ^21,35^.

### CD73^+^CD80^+^ RR-specific memory B cells form short-lived plasmablasts after subsequent challenge

Although RR-specific, class-switched CD73^+^ CD80^+^ MBCs could form after a *Plasmodium* infection or immunization in a GC-independent manner (Figs. 1–2), it was unclear how these MBCs would respond to antigen re-exposure. We therefore infected C57BL/6 mice with 2000 *Py^Pf^*^CSP^ sporozoites and allowed memory populations to form as described above (Fig. 3a). At least 60 days after infection, we rechallenged previously infected mice with 25*μ*g RT-I53-50 and tracked B cell responses up to 28 days later (Fig. 3a). We detected a rapid expansion of RR- specific B cells as early as three days post-challenge (Fig. 3b). These cells were largely short-lived PBs that contracted by day 14 (Fig. 3b,c). This rechallenge-induced PB response was faster than the PB response seen after a primary RT-I53-50 immunization (Fig. 3b compared to Fig. 2a), demonstrating MBC derivation, rather than activation of naïve B cells. In contrast, immunization induced little to no expansion of CT-specific B cells in previously infected mice and very few CT-specific PBs (Fig. 3b, c). We also measured epitope-specific serum antibody responses and detected an increase in RR-specific serum IgG antibody concentrations at day 7 post rechallenge, immediately after the PB burst, but no significant change in CT-specific antibody titers (Fig. 3d). CD73^+^CD80^+^ RR-specific MBCs are therefore functional, forming a rapid, antibody-secreting PB response after secondary antigen exposure, while CT-specific secondary responses are minimal, again highlighting the different responses to these two different regions in the CSP antigen.

**Fig. 3:**
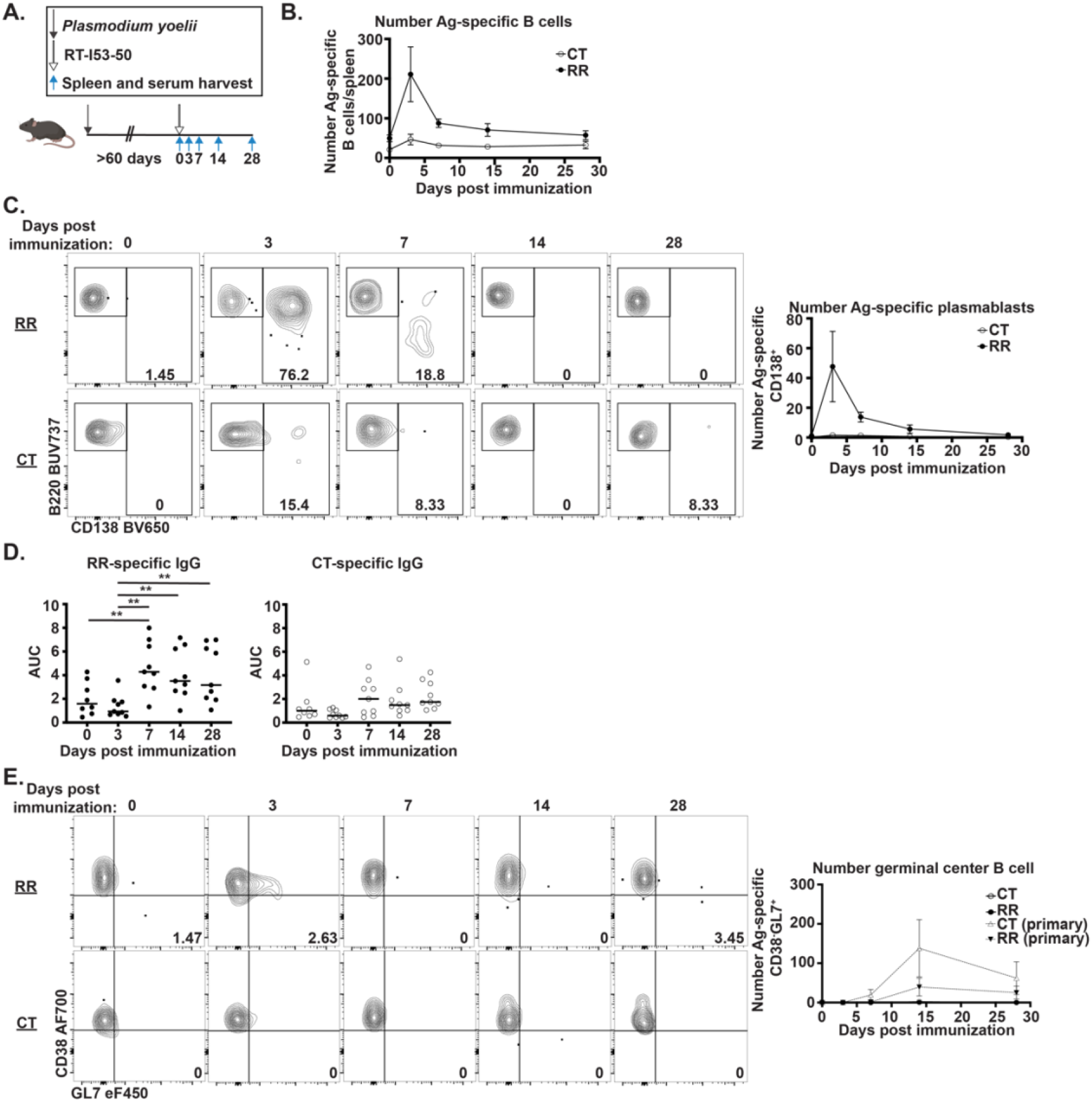
RR-specific memory B cells form an optimal PB response upon rechallenge, but do not form secondary GCs. **a,** C57BL/6 mice were infected with 2000 *Py^Pf^*^CSP^. At least 60 days later, mice were immunized with 25*µ*g RT-I53-50. Serum and spleens were harvested from mice prior to immunization or indicated days post immunization. Splenic RR- and CT-specific B cells were isolated and phenotyped. **b,** Total numbers of RR-and CT-specific B cells after immunization. **c,** Formation of RR- and CT-specific PBs (CD138^+^) at indicated times post-immunization. **d,** RR-or CT-specific serum IgG at indicated times post-immunization. **e,** RR- and CT-specific GCs (CD38^-^GL7^+^) at indicated times post-immunization. GC B cells after a primary RT-I53-50 immunization of previously naïve mice are shown as a control (primary infection, dotted lines). n = 8-9. Statistical analysis by Mann-Whitney **p<0.01

### RR-specific MBCs do not enter secondary GCs or proliferate into new, long-lived memory after challenge

Our data demonstrate that wild-type *Plasmodium* infection predominantly drives a GC-independent CD80^+^CD73^+^ RR-specific MBC population (Figs. 1–2) that can rapidly form functional PBs after challenge (Fig. 3b-d), yet perhaps a lack of GC-experience altered the MBCs in other ways that could lead to waning immunity. A lack of maintenance of these GC-independent B cells could provide an alternative mechanism underlying the loss of protection over time that has eluded our understanding over the past decades. We reasoned that this could be influenced by the demonstrated ability of CD80^+^CD73^+^ MBCs to form PBs at the expense of re-entering GCs^27,29,36^. To understand whether new memory populations are generated upon secondary antigen exposure, we first looked at the formation of secondary GCs after a rechallenge (Fig. 3a). After RT-I53-50 challenge, we observed no CD38^-^GL7^+^ RR- or CT-specific GC B cells at any time point (Fig. 3e). This contrasts with a primary RT-I53-50 immunization that generates robust CT- specific GCs and some RR-specific GCs (Fig. 3e control dotted grey line, Fig. 2c). These data suggest that having preestablished RR-specific memory after *Plasmodium* infection prevents the formation of secondary GCs, likely due to the terminal differentiation of CD80^+^CD73^+^ MBCs that predominantly form short-lived PBs.

It was also possible that the MBC population can be renewed by existing MBC proliferation independent of a GC. To assess whether the RR-specific MBCs we characterized in response to a primary wild-type *Plasmodium* infection could proliferate to form new long-lived MBCs after challenge, we again infected mice with 2000 *Py^Pf^*^CSP^ sporozoites, allowed memory populations to develop, and rechallenged the mice with 25*μ*g RT-I53-50 (Extended Data Fig. 5a). However, this time we also provided daily treatment of BrdU during the expansion of the RR-specific MBCs present (days 1-6, Fig. 3b) to label any proliferating cells (Extended Data Fig. 5a). We then assessed BrdU incorporation in RR-specific PBs, naïve IgD B cells, and MBCs at 3- or 14-days post immunization (Extended Data Fig. 5a). As expected, 3 days post immunization essentially all the RR-specific PBs had BrdU incorporation, demonstrating their rapid proliferation from quiescent MBCs (Extended Data Figs. 5b,c). Three days post RT-I53-50 boost, most RR-specific MBCs were BrdU^+^, indicating that they were replicating rapidly (Extended Data Figs. 5b,c). However, by day 14 almost all BrdU^+^ RR-specific MBCs had disappeared, leaving only BrdU^-^ populations (Extended Data Figs. 5b,c). This suggests that the RR-specific MBCs responding to challenge rapidly replicate into short-lived effectors (as indicated by BrdU^+^ PBs 3 days post challenge), but do not survive until later timepoints. Overall, these data support a model where the RR-specific MBCs generated after *Plasmodium* infection are poised to differentiate into functional, but short-lived, PBs upon recall rather than renewing durable memory populations.

### Multiple *Plasmodium* infections results in the depletion of RR-specific memory B cells

The differentiation of MBCs into short-lived PB upon rechallenge (Fig. 3c) may be temporarily beneficial and is associated with protection in clinical trials^4,37,38^, but could have negative long-term consequences if long-lived MBCs are not replenished. We hypothesized that differentiation into short-lived effectors may therefore eventually lead to a loss of functional memory after repeated infection. To test this hypothesis, we infected C57BL/6 mice either once, twice, three times, or four times with 2000 *Py^Pf^*^CSP^ sporozoites, waiting at least 28 days between each infection (Fig. 4a). We quantified the number of RR-specific B cells and the number of class-switched and/or CD73 and CD80 expressing RR-specific B cells one month after the final infection (which ranged from 1-4 infections). We found a significant decrease in the total number of RR-specific B cells and the number of class-switched B cells as early as after the second infection that remained low after all additional infections (Figs. 4b,c). Similarly, we detected a decrease in the number of CD73 and CD80 expressing MBCs (Figs. 4b,c). The number of class-switched and CD73^+^ CD80^+^ MBCs dropped dramatically after four infections (Figs. 4b,c). These data provide strong evidence that after repeated infections there is a loss of functional memory due to a lack of maintenance of this important population and, for the first time, suggests a mechanism explaining the short-lived nature of malaria vaccines.

**Fig. 4:**
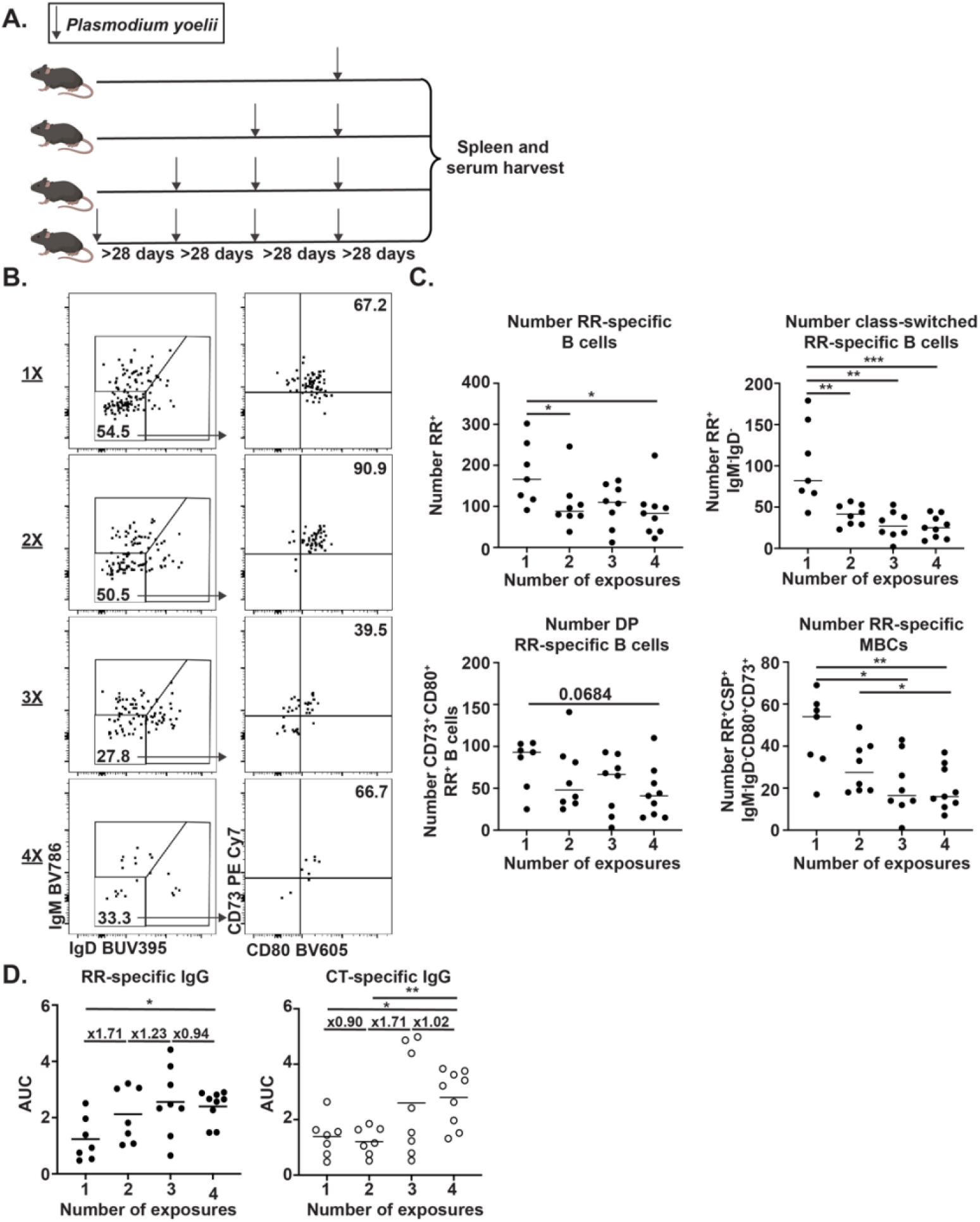
Multiple *Plasmodium* infections result in a depletion of RR-specific memory. **a,** C57BL/6 mice were infected with 2000 *Py^Pf^*^CSP^ up to four times with at least 28 days between infections. Serum and spleens were harvested at least 28 days after the last infection. RR-specific B cells were isolated and phenotyped. **b,** Expression of CD73, CD80, and isotype of RR-specific B cells after 1-4 infections. **c,** Quantification of B, total number of RR-specific B cells and number of class-switched (IgD^-^IgM^-^) and/or CD73^+^CD80^+^ RR-specific B cells. **d,** RR- or CT-specific serum IgG antibody at least 28 days after indicated number of infections. n = 7-9. Statistical analysis by Mann-Whitney *p<0.05, **p<0.005, ***p<0.001

Because we detected a loss of RR-specific MBCs, we next assessed whether additional infections would likewise impact IgG antibody titers. Mice that had been infected only once had low levels of RR-specific IgG 28 days later, consistent with data showing that primary infection mostly generates IgM PBs (Fig. 4d and Extended Data Fig. 2). When mice were infected with *Py^Pf^*^CSP^ a second time, there was a 70% increase in RR-specific IgG antibody (Fig. 4d), like that detected in Fig. 3d, that was not associated with an increased MBC response (Fig. 4b,c). However, additional infections gradually lost the ability to boost RR-specific IgG serum antibody levels (Fig. 4d). A third infection only increased titers 23% over the previous levels, and a fourth infection failed to induce a boost at all. This is consistent with recent data in clinical trials that continuous immunization boosts eventually stop generating increased RR-specific antibody titers^6,39^.

Interestingly, when we assessed CT-specific antibody titers at the same timepoints we detected an increased amount of CT-specific IgG antibody by the fourth infection (Fig. 4d). It is possible that with the loss of RR-specific responses, CT-specific B cells may gain access to antigen or T cell help and enable a CT-specific response. The role of B cell responses towards the CT on protection from *Plasmodium* are incompletely understood, but there is evidence that CT-specific antibodies bind sporozoites poorly and provide little protection in preclinical models^40,41^. Consistent with these data, a co-submitted manuscript (Netland et al.) demonstrates using a longitudinal analysis in human PBMCs that while RR-specific B cells dominate early responses, long-term memory primarily comprises CT-specific B cells that are poorly protective in a murine model of infection. It is possible that the differentiation of RR-specific MBCs exclusively into PBs may not only be depleting protective RR-specific memory but may also be permitting the expansion of humoral responses towards less protective epitopes.

### RR valency and density influence B cell differentiation and contribute to impaired memory formation

Our data indicate that RR-specific MBCs are impaired in their ability to replenish the MBC pool which leads to their depletion after multiple antigen exposures (Fig. 4). We next wanted to understand what leads a B cell down this dysfunctional trajectory. We hypothesized that the high RR valency and density on the sporozoite surface contribute to this phenotype. To test how RR antigen valency and density influence B cell differentiation, we generated modified RT-I53-50 NPs that contain a range of repeat valency and density. Some particles contain 4 NANP repeats (NANP_4_, low valency) whereas others contain 19 NANP repeats (NANP_19_, high valency) (Fig. 5a). Additionally, we generated NPs with 30 or 60 copies of NANP_4_ or NANP_19_ on the surface, representing low or high antigen density, respectively (Fig. 5a). We immunized mice with equal concentrations of RR antigen displayed by these four NPs and assessed responses seven days later (Fig. 5b). We found higher valency or density led B cells to differentiate into PBs at a higher frequency (Figs. 5c,d), consistent with antigen strength influencing B cell differentiation^42^. In contrast, lower valency or density NPs drove formation of RR-specific GCs that persisted up to 30 days post immunization, whereas we were largely unable to detect any GCs after immunization with antigen high valency and density (Fig. 5d). Importantly, this had consequences on memory formation, as high antigen valency and density led to significantly fewer persisting RR-specific class-switched, CD73^+^ CD80^+^ MBCs (Fig. 5e). Overall, these data demonstrate that the high antigen valency and density of the RR likely contribute to impaired B cell responses.

**Fig. 5:**
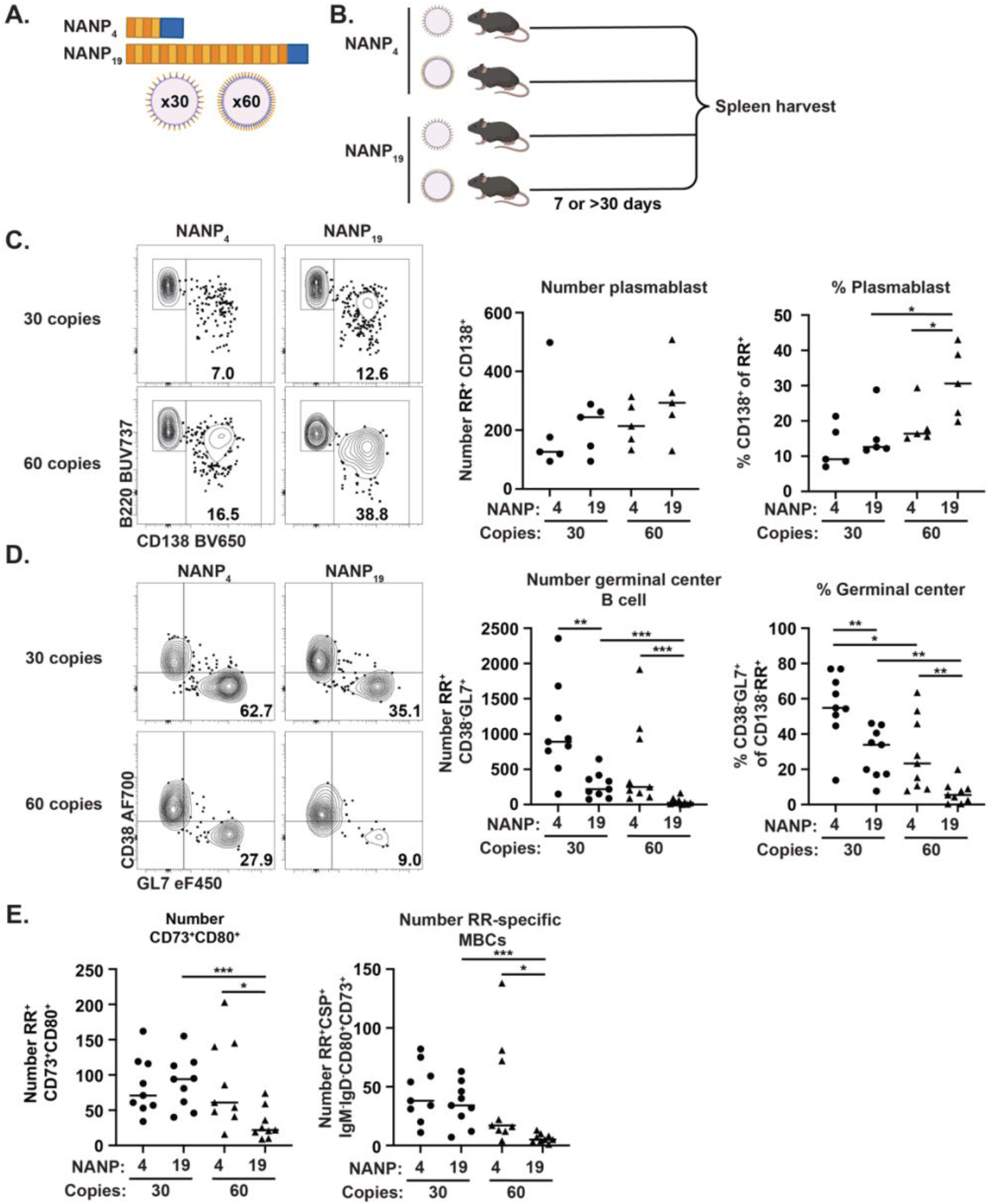
Lower RR valency or density improves acute and memory B cell responses. **a,** NPs containing either 30 or 60 copies of NANP_4_ or NANP_19_ were generated. **b,** C57BL/6 mice were immunized with 0.63*µ*g NANP antigen and spleens were harvested 7 or at least 30 days later. Splenic RR-specific B cells were isolated and phenotyped. **c,** Formation of RR-specific PBs (CD138^+^) at 7 days post-immunization. **d,** RR-specific GCs (CD38^-^GL7^+^) at least 30 days post-immunization. **e,** Number of CD73^+^CD80^+^ (left) and CD73^+^CD80^+^IgM^-^IgD^-^ (right) RR- specific B cells at least 30 days post-immunization. n = 5-9. Statistical analysis by Mann-Whitney *p<0.05, **p<0.005, ***p<0.001

### Vaccination of children in endemic areas induces RR-and CT-specific B cell responses

While there were distinct phenotypic differences of RR-and CT-specific B cells in our murine model, it was also important to determine if these findings translated to clinical samples. To do so, we first assessed the number and phenotype and of RR- and CT-specific B cells in PBMCs from donors participating in the Bagamoyo Sporozoite Vaccine trial 2 (BSPZV2^43^, clinicaltrials.gov identifier NCT02613520) trial at the Ifakara Health Institute in Tanzania. In this trial, children under 18 years old were immunized with three doses of irradiated sporozoites intravenously and PBMCs were collected at the time of the first vaccination and throughout the course of immunization (Fig. 6a). We assessed the number and phenotype of RR-specific B cells and found that children had very few RR-specific B cells prior to vaccination (V1), but vaccination induced a robust expansion of RR-specific B cells by day 14 after the first immunization (V1+14, Fig. 6b, Extended Data Fig. 6). To learn more about the distinct features of the RR-specific MBCs, we next assessed isotype and expression of cell surface proteins associated with populations of MBCs in humans. We detected an expansion of antigen-experienced (CD21^-^, CD27^-/+^) and IgG-expressing RR-specific B cells after each immunization (Fig. 6c). Overall, approximately 70% of children responded to vaccination, increasing the number of total RR-specific B cells as well as RR-specific MBCs and IgG B cells (Fig. 6d right). This response rate was detected as early as 14 days post the first vaccine (V1+14, Fig. 6d left) and was maintained through the immunization course (Fig. 6d right). Overall, these data indicate that vaccination of children expands RR-specific responses and specifically MBCs after each vaccination.

**Fig. 6:**
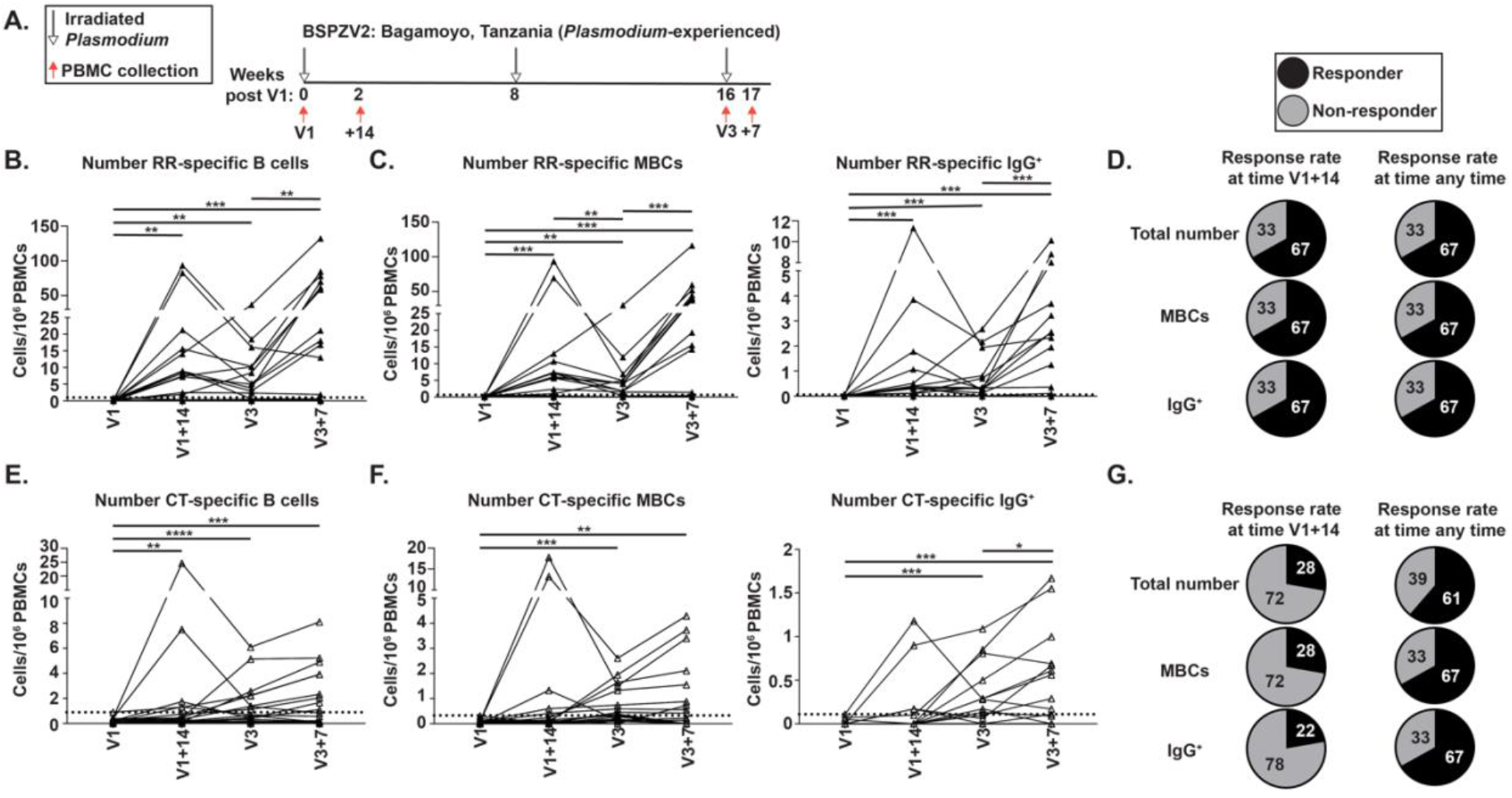
Vaccination of children in endemic areas induces RR-and CT- specific B cell responses. **a,** Schematic of BSPZV2 clinical trials for children donors. **b,** The total number of RR-specific B cells at the indicated timepoints **c,** The total number of RR-specific MBCs (CD21^-^CD27^+/-^, left) or RR-specific IgG B cells (right) at the indicated timepoints. **d,** The RR-specific response rate at early timepoints (V1+14, left) or at any timepoint (right) **e,** The total number of CT-specific B cells at the indicated timepoints. **f,** The total number of CT-specific MBCs (CD21^-^CD27^+/-^, left) or CT-specific IgG B cells (right) at the indicated timepoints. **g,** The CT-specific response rate at early timepoints (V1+14, left) or at any timepoint (right). The dotted line represents three standard deviations above the average at time V1. Donors reaching values above that line were designated as responders (black, pie charts) and those below as non-responders (grey, pie charts). n = 18 people per time point. Statistical analysis by Wilcoxon matched-pair signed rank test *p<0.05, **p<0.005, ***p<0.001, ****p<0.0001

We similarly assessed whether vaccination would generate a CT-specific response and found that vaccination induced an expansion of CT-specific cells in terms of total number, antigen-experienced, and IgG-expressing cells (Fig. 6e,f). Importantly, whereas significant increases in RR-specific responses occurred as early as the first immunization, the expansion of CT-specific B cells largely occurred after later vaccinations. While approximately 70% of children had increases in RR-specific responses by V1+14 (Fig 6d left), only 30% of children had CT-specific responses at this time (Fig. 6g left). Additional immunizations eventually enhanced the CT-specific response to the level of RR-specific response rate (70%, Fig. 6g right). Consistent with our murine data (Fig. 4d), previous published work^44^, and data from an additional clinical trial analyzed in a co-submitted manuscript (Netland et al.), these data suggest that the immunodominant RR monopolizes early responses, but responses to low avidity domains may be generated after repeated antigen exposure. In sum, these data indicate that a multiple vaccination series in children generates RR-and CT-specific responses in children, though with different kinetics.

### Adults in malaria-endemic regions have impaired RR-and CT-specific B cell responses after vaccination

Our data in a mouse model suggest that RR-specific MBCs are depleted after repetitive *Plasmodium* infection due to inefficient renewal of memory. As individuals in malaria endemic areas experience up to an estimated 300 infectious mosquito bites a month in a rainy season^45,46^, we hypothesized that adults would have an impaired RR-specific response compared to children. To test how ongoing *Plasmodium* exposure impacts the response to vaccination, we next assessed RR-and CT-specific B cells in adult donors from the same BSPZV2 trial (Fig. 7a). To control for age differences, we also assessed responses in a second trial that immunized *Plasmodium*-naïve adults, (IMRAS^47^, clinicaltrials.gov identifier NCT01994525). Donors in this trial were given five immunizations of ∼200 bites from radiation attenuated sporozoite-infected mosquitoes (Fig. 7a).

**Fig. 7:**
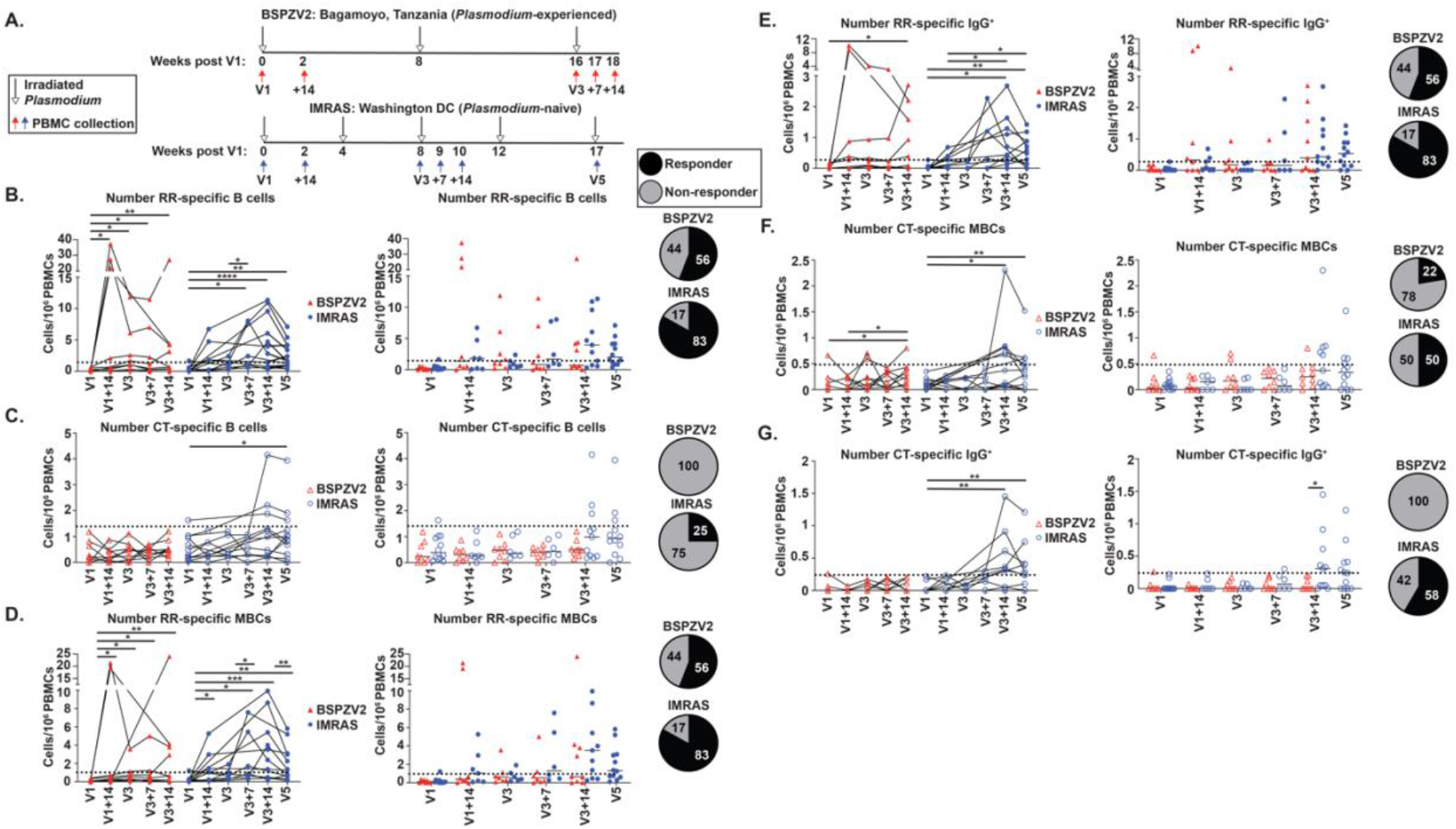
Adults living in endemic areas have impaired RR- and CT-specific responses to CSP vaccination. **a,** Schematic of *Plasmodium*-naïve (IMRAS, blue) and *Plasmodium*-experienced (BSPZV2, red) clinical trials for adult donors. **b,** The total number of RR-specific B cells at indicated timepoints. **c,** The total number of CT-specific B cells at indicated timepoints. **d-e** The number of RR-specific MBCs (CD21^-^CD27^+/-^, **d**) or IgG B cells (**e**) at indicated timepoints. **f-g,** The number of CT-specific MBCs (CD21^-^CD27^+/-^, **f**) or IgG B cells (**g**) at indicated timepoints. The dotted line represents three standard deviations above the average at time V1. Donors reaching values above that line at any timepoint were designated as responders (black, pie chart) and those below as non-responders (grey, pie chart). n = 6-12 people per time point. Statistical analysis by Wilcoxon matched-pair signed rank test for longitudinal analysis and multiple unpaired Mann-Whitney tests for group analysis. *p<0.05, **p<0.005, ***p<0.001, ****p<0.0001

We assessed the number of RR- and CT-specific B cells in *Plasmodium*-experienced and -naïve adults prior to the first immunization. As donors who live in endemic areas had likely been infected with *Plasmodium* repeatedly throughout their lives^45,46^, we expected some numerical advantage compared to donors who had never been exposed to CSP. Strikingly, however, we observed no difference in the number of RR-specific B cells at the time of initial vaccination (V1) in previously malaria-exposed adults compared to malaria-naïve people (Fig. 7b). We suspect that the lack of memory is driven by the RR-specific memory depletion after multiple infections, as demonstrated in our mouse model (Fig. 4). We similarly found no difference in the number of CT-specific B cells at the time of V1 (Fig. 7c), consistent with our murine data suggesting that infection does not generate a robust CT-specific memory population.

We next assessed whether the number of RR- and CT-specific B cells could be enhanced with vaccination. We found that only 4 out of 9 *Plasmodium*-experienced adults exhibited an expansion of RR-specific B cells at any timepoint (Fig. 7b). In contrast, as quickly as two weeks post the first immunization (V1+14), 10 out of 12 *Plasmodium*-naïve adults had an expansion of RR-specific B cells (Fig. 7b), similar to the response rate seen in children living in endemic areas (Fig. 6b). This suggests that having prior *Plasmodium* experience impairs the RR-specific response to vaccination. When we assessed changes in the number of CT-specific B cells, we found that *Plasmodium*-experienced adults completely failed to induce any expansion of CT-specific B cells throughout the entire course of immunization whereas *Plasmodium*-naïve individuals had a slight increase in CT-specific B cell number that was only detectable at later timepoints after the third vaccination (V3+14 and V5, Fig. 7c).

Similarly, at the time of the first vaccination (V1) we found that *Plasmodium*-experienced adults did not have elevated levels of RR-specific antigen-experienced B cells (CD21^-^, CD27^-/+^) or IgG B cells compared to a *Plasmodium*-naïve donor (Figs. 7d,e). This again suggests that prior infection is not generating durable, RR-specific memory. After vaccination, 83% of previously naïve donors had a detectable increase in the number of IgG and antigen-experienced RR- specific B cells, whereas almost half of the *Plasmodium*-experienced adults did not have any increase in the number of these cells, with the exception of a few outliers that generated a response that more resembled malaria-naïve donors (Figs. 7d,e). Further, only *Plasmodium*-naïve individuals and no *Plasmodium*-experienced adults had a detectable increase in antigen-experienced, IgG^+^ CT-specific B cells (Figs. 7f,g). Like we detected in children in the BSPZV2 trial, the CT-specific response was again only detected at later timepoints after vaccination 3, (V3+14 and V5) compared to the quick response of RR-specific B cells (Figs. 7f,g).

Overall, these data indicate that ongoing exposure and parasite-driven immunomodulation negatively impact the B cell response to vaccination. Individuals with fewer prior exposures (*Plasmodium*-naïve adults or children in living in endemic regions) generate superior expansion and differentiation of CSP-specific B cells compared to adults in endemic areas. This suggests that vaccinating children as young as possible is optimal to maximize vaccine efficacy, while vaccinating adults with significant prior infection may not enhance humoral immunity towards sporozoite antigens.

## Discussion

Though CSP-based vaccines have reduced rates of hospitalization and deaths due to malaria^2^, efficacy of these vaccines is short-lived and inferior in *Plasmodium*-experienced individuals^3–6,9^. We sought to understand the cellular mechanisms that contribute to these defects. Using both murine samples and human PBMCs, we have demonstrated for the first time that the repetitive nature of the RR-specific contributes to the generation of dysfunctional MBCs that are poised to differentiate into short-lived effectors at the expense of generating a long-lived, repopulating memory. This eventually leads to a loss of memory after repeated infections. This depletion has severe consequences on current vaccination strategies and has implications on the development of the next generation of CSP-based vaccines.

This work leads to many new questions, most importantly understanding how the repetitive nature of the RR-specific B cells influences MBCs differentiation such that they only generate short-lived effectors upon recall. Our data suggest that the repetitive nature of the epitope itself contributes to the dysfunctional B cell differentiation. We demonstrate that higher antigen valency or density polarizes RR-specific B cells towards short-lived PBs and not GC B cells during a primary response. In line with this hypothesis, previous research has demonstrated that other vaccines containing repetitive high avidity epitopes, such as the 23-valent pneumococcal polysaccharide vaccine, leads to significant B cell receptor (BCR) crosslinking and differences in signaling^24^. This induces a short-lived PB response and depletes long-lived memory. Though the mechanism may be slightly different (as the response to the pneumococcal vaccine is T-independent whereas RR- specific responses depend on T cells (Extended Data Fig. 4)^24,48^), the phenotypes mimic the short-lived effector response we detect in our studies. It is possible that *Plasmodium* spp. evolved this repetitive protein as a strategy to influence B cell differentiation by excessive BCR crosslinking, skewing these cells towards short-lived effectors and disrupting the ability to replenish memory after repeated infection.

If the repetitive nature of the RR contributes to the aberrant MBC polarization, use of vaccines that integrate the RR into the platform, like RTS,S, will not generate long-lived immunity in response to repeat exposure. Our data suggest that, in contrast, vaccine boosters may impair immunity further by depleting functional RR-specific MBCs. In line with this hypothesis, anti-CSP antibody titers generated from an additional RTS,S booster have already been demonstrated to only induce serum antibody titers half as large as the initial vaccination^4,6^. Consistent with our data, this phenotype is domain-specific, as RR-specific antibody titers not only wane more rapidly than CT-specific titers but exhibit a diminished boosting response^39^. Whereas booster vaccination increases CT-specific titers to levels exceeding those achieved after primary immunizations, RR- specific titers remain below their post-prime levels even after boosting^39^. Furthermore, data from a co-submitted manuscript (Netland et al.) conclusively demonstrate on a cellular level that in *Plasmodium*-naïve individuals, multiple boosts with CSP-based vaccines eventually leads to a loss of RR-specific memory at late timepoints. This suggests that inclusion of the repetitive antigen itself contributes to the aberrant memory polarization. Vaccines that depend upon a protective, repetitive epitope like the newly modified RTS,S vaccine, called R21/Matrix-M (recently introduced into the clinic) should be further analyzed for similar longevity issues^1,49^. Both RTS,S and R21 utilize the same virus-like particle scaffold, but R21 contains approximately five times as much CSP on the surface^49^. We suspect that integration of additional RR epitopes may enhance early antibody production but further skew MBCs towards a short-lived phenotype and away from differentiation programs associated with long-lived memory. This could result in a subsequent reduction of vaccine durability.

In our studies, we determined that RR-specific MBCs were not proliferating into new, long-lived memory. However, new memory can also form when naïve B cells encounter antigen during a second infection. We were able to detect IgD-expressing RR-specific B cells even after four *Plasmodium* infections (Fig. 4b), presenting the question of why the naïve cells were not participating in the response after additional antigen exposures. Recent studies have demonstrated that antibody feedback prevents generation of RR-specific responses after vaccination boosts^44,50^. As we detect a large PB burst and RR-specific antibody present at the time of recall (Fig. 3c,d and Fig. 4d), the naïve RR-specific B cells are potentially not gaining access to antigen by this mechanism. The combination of naïve cells being unable to participate in a reaction and aberrant memory populations would lead to the loss of RR-specific memory we detect in our studies.

A major motivation for this research was to identify mechanisms to improve malaria vaccines. Our data demonstrate that children have more robust RR-specific B cell responses (Fig. 6), likely due to fewer prior infections, highlighting this is still a viable strategy for young children but not necessarily beyond this age group in malaria endemic regions. Furthermore, immunizing children with vaccines containing fewer repetitive segments may be an effective strategy to limit excessive BCR crosslinking. However, these strategies still may not work long-term if *Plasmodium* infection after vaccination disrupts RR-specific memory due to the many repeats in the parasite itself. More work on the impact of infection after immunization is needed to elucidate these details. Alternatively, vaccines generating long-lived plasma cells (LLPCs) that produce RR-specific antibody would be a superior platform. LLPCs are largely refractory to secondary antigen exposures as they do not express much BCR, but rather consistently produce antibody^51^. If we could generate vaccines that maintain a baseline RR-specific antibody titer that cannot be disrupted with additional antigen exposures, this could provide long-lived immunity regardless of subsequent infection.

In conclusion, we have identified cellular mechanisms that explain the impaired longevity of protective RR-specific humoral immunity, especially in *Plasmodium*-experienced individuals. Our data provide compelling evidence that RR-specific memory is depleted with continuous antigenic exposure and especially in response to infection in both murine models and human PBMCs. These data have important consequences for future vaccine design, particularly when developing vaccines targeting individuals living in endemic areas.

## Methods

### Protein expression and tetramer generation

NANP6 (RR, NANPNANPNANPNANPNANPNANP) or *Pf*16 (CT, EPSDKHIKEYLNKIQNSLSTE WSPCSVTCGNGIQVRIKPGSANKPKDELDYANDIEKKICKMEKCS) peptides containing an N-terminus biotin were purchased from Biomatik. Recombinant full-length *Pf*CSP was expressed using HEK293F cells (Thermo) and then column purified using the Ni-NTA kit (Millipore Sigma). *Pf*CSP was then biotinylated using the EZ-Link Sulfo-NHS-LC-Biotin kit (Thermo) according to the manufacturer’s instructions. Biotinylated *Py^Pf^*^CSP^, NANP6, and *Pf*16 tetramers reagents were generated as previously described^52^. Briefly, biotinylated proteins were incubated with streptavidin-PE, AF488, or APC at room temperature for 30 minutes. The tetramer fraction was centrifuged in a 100KD (*Py^Pf^*^CSP^) or 30KD (NANP6 and *Pf*16) amicon column (Millipore) to filter out non-tetramerized proteins. Probe specificity was validated by sorting tetramer positive cells, sequencing and expressing BCRs, and performing RR- or CT-specific ELISAs (Netland et al.).

### Nanoparticle production and purification

Protein nanoparticle used in this study was based off the two-component icosahedral platform, I53-50^33^. RT-I53-50A, which contains 19 NANP repeats and the CT domain of CSP, was codon-optimized in pET29b+ for expression in *E. coli* as previously described^34^ and transformed into BL21(DE3) *E. coli* (New England Biosciences, Massachusetts, USA). Transformed colonies were used to inoculate LB containing 50 *μ*g/mL kanamycin and cultures were grown overnight in a shaking incubator at 37°C and 200 RPM. The following morning, 500 mL of ZY Autoinduction medium (ZY media, 50XM Salts, 50X5052, 1M MgSO4, and 1000X Trace metals) with 50 µg/mL of kanamycin in a baffled 2L flask was inoculated with 5-10 mL of overnight culture. Cultures were grown shaking at 37°C for two hours and temperature was reduced to 18°C for overnight expression. Overnight cultures were harvested by centrifugation at 4000 rcf at 4°C for 20 min.

The cell pellet was resuspended in a buffer containing 50 mM Tris, 500 mM NaCl, 1 mM DTT, 30 mM imidazole, 1mM PMSF, 10 µg/mL DNase, pH 8.0. Re-suspended cells were homogenized and lysed through a Microfluidizer (M-110P, Microfluidics, USA) and clarified by centrifugation to separate insoluble cell debris from supernatant. The resulting supernatant was purified using Ni-NTA resin (Qiagen, Venlo, Netherlands) that was pre-equilibrated in wash buffer (50 mM Tris, 500 mM NaCl, 30 mM imidazole, pH 8.0), washed with five column volumes of wash buffer, and eluted with buffer containing 50 mM Tris, 500 mM NaCl, 300 mM imdazole pH 8.0. The resulting protein eluate was concentrated and purified by size exclusion chromatography (SEC) on a Superdex 200 Increase 10/300 GL column (Cytiva) in a buffer containing 50 mM Tris, 500 mM NaCl pH 8.0. Peak fractions corresponding to the correct molecular weight were collected and purity was verified by SDS-PAGE.

For high density immunogens, nanoparticles were assembled by mixing purified full-length (NANP_19_) RT-I53-50A or I53-50A components displaying truncated RR (NANP_4_) with I53-50B component in a 1.1:1 ratio as previously described^53^. To generate particles corresponding to 30 NANP copies, bare I53-50A (lacking CSP) was mixed at a 1:1 molar ratio with either NANP_4_- or NANP_19_-expressing I53-50A. Because RT-I53-50 NPs can display a maximum of 60 CSP antigen copies, incorporation of bare I53-50A at equal molar ratios reduces the antigen display capacity in half, resulting in 30 CSP antigen copies. This mixture was then incubated with I53-50B as above to generate low density particles. The nanoparticle assembly was purified by SEC in 50 mM Tris, 500 mM NaCl pH 8.0 buffer using a Superose 6 Increase 10/300 GL column (Cytiva) to remove residual components. Fractions corresponding to the expected size of the nanoparticle assembly were collected and pooled, and concentration was determined using a UV-vis spectrophotometer (Agilent Cary 8454). Nanoparticle was further characterized by dynamic light scattering (DLS) to confirm the expected hydrodynamic diameter and monodispersity, and negative-stain electron microscopy (nsEM) was carried out to confirm the presence of nanoparticle with the expected icosahedral morphology. Lastly, endotoxin measurements were taken to ensure an EU below 100 EU/mg. Nanoparticle was snap frozen and stored at -80°C before further use in animal experiments.

### Parasite growth and sporozoite isolation

Swiss Webster or C57BL/6 mice were injected with blood infected with *Py^Pf^*^CSP^ ^25^ to begin the growth cycle and used to feed female *A. stephensi* mosquitos after gametocyte exflagellation. Salivary gland sporozoites were isolated on days 13-17 post infectious blood meal. For indicated experiments, sporozoites were exposed to 200 gray irradiation.

### Mice and parasite infections and treatments

7-12-week-old C57BL/6, CD4cre^+^, TCRα^-/-^, Bcl6^fl/fl^ mice were purchased from Jackson Laboratory and then bred and maintained under specific pathogen free conditions at the University of Washington. CD4cre heterozygotes were bred to Bcl6^fl/fl^ mice to generate CD4cre^+^Bcl6^fl/fl^ and CD4cre^-^Bcl6^fl/fl^ strains. Genotypes were confirmed by PCR using the primers listed in Table 1. For parasite infections, mice were anaesthetized via isoflurane (Piramal) and injected with 2000 *Py^Pf^*^CSP^ sporozoites or 50000 irradiated *Py^Pf^*^CSP^ sporozoites in 200*μ*L PBS intravenously via retro-orbital injection. One to two weeks after infection, parasitemia by was confirmed by detection of Hoeschst^+^ red blood cells via flow cytometry. For RT-I53-50 immunizations, 25*μ*g total protein (8.3*μ*g CSP antigen) was diluted in 2X Sigma Adjuvant System. Mice were immunized with 100*μ*L RT-I53-50/1X SAS solution by intraperitoneal injection. For varied RR valency and density NP experiments, mice were immunized with 0.63*μ*g NANP antigen in 1X Sigma Adjuvant System by intraperitoneal injection. Mice that received BrdU treatment were dosed with 1mg BrdU intraperitoneally (Sigma, 100-200*μ*L volume, diluted in PBS) one hour prior to RT-I53-50 immunization. Mice were treated with additional 1mg BrdU daily until harvest. All animal experiments were ethically performed in accordance with the University of Washington Institutional Animal Care and Use Committee guidelines.

**Table 1:**
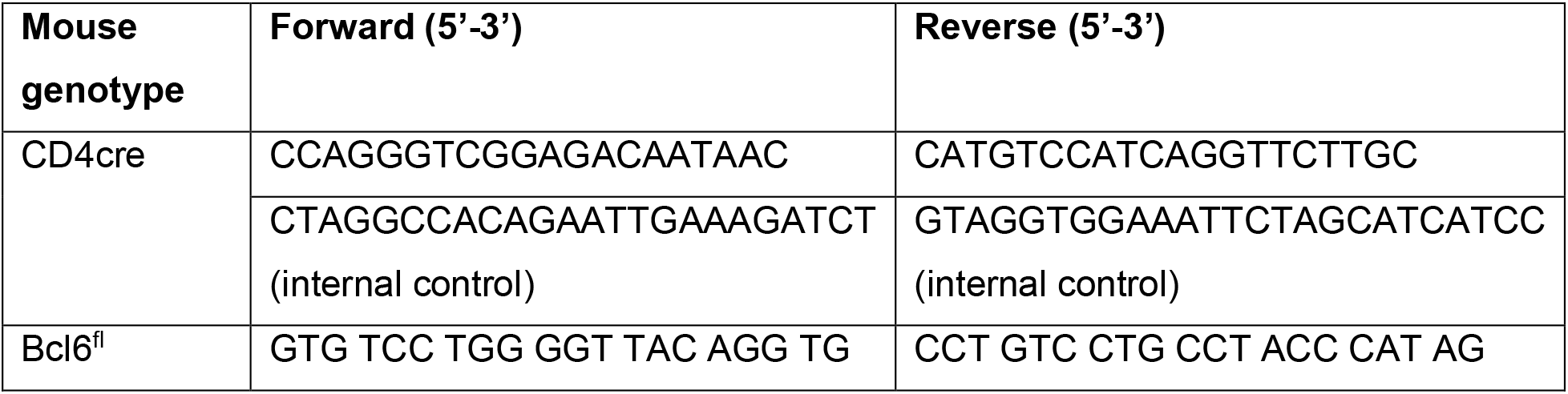
Genotyping primers.

### Cell isolation, cell enrichment, and flow cytometry

For murine samples, mice were euthanized with CO_2_ asphyxiation and the spleens were mashed with the back of a 3cc syringe through Nitex mesh (Amazon.com) to generate single cell suspensions. For human samples, PBMCs were thawed at 37°C and washed twice in FACS buffer. Human and murine single cell suspensions were stained with PE-AF594-AF650 decoy reagent for 15 minutes at room temperature, followed by 30 minutes at 4°C in *Pf*CSP-PE, NANP6-AF488, and *Pf*16-APC tetramer. Cells were then washed and incubated with anti-PE microbeads (Miltenyi Biotech) for 20 minutes at 4°C. *Pf*CSP-specific B cells were then enriched as previously described^52^. Enriched cell fraction was incubated with Fc block for 15 minutes at 4°C followed by incubation with surface markers on ice for 30 minutes with antibodies listed in Table 2. For samples treated with BrdU, integration of BrdU into genomic material was assessed by per the manufacturer’s protocol (Biolegend). All cells were run on Symphony and analyzed using FlowJo software.

**Table 2:** Flow cytometry reagents.

| **Antibody** | **Clone** | **Color** | **Source** |
| --- | --- | --- | --- |
| anti-human CD20 | 2H7 | PerCP-Cy5.5 | BD Biosciences |
| anti-human CD38 | HIT2 | AF700 | Biolegend |
| anti-human IgA | IS11-8E10 | biotin | Miltenyi Biotec |
| Streptavidin |  | BUV395 | BD Biosciences |
| anti-human CD27 | M-T271 | BV421 | Biolegend |
| anti-human IgM | MHM-88 | BV510 | Biolegend |
| anti-human CD21 | HB5 | SB600 | Invitrogen |
| anti-human CD14 | MφP9 | BV711 | BD Biosciences |
| anti-human CD16 | 3G8 | BV711 | BD Biosciences |
| anti-human CD3 | UCHT1 | BV711 | BD Biosciences |
| anti-human IgG | G18-145 | BV786 | BD Biosciences |
| anti-human IgD | IA6-2 | PE-Cy7 | BD Biosciences |
| anti-human CD19 | SJ25C1 | APC-Cy7 | BD Biosciences |
| anti-mouse B220 | RA3-6B2 | BUV737 | BD Biosciences |
| anti-mouse CD4 | GK1.5 | BUV805 | BD Biosciences |
| anti-mouse CD138 | 281-1 | BV650 | BD Biosciences |
| anti-mouse CD38 | 90 | AF700 | Invitrogen |
| anti-mouse GL7 | GL7 | eF450 | Invitrogen |
| anti-mouse IgM | II/41 | BV786 | BD Biosciences |
| anti-mouse IgD | 11-26c.2a | BUV395 | Biolegend |
| anti-mouse CD73 | eBioTY/11.8 | PE-Cy7 | Invitrogen |
| anti-mouse CD80 | 16-10A1 | BV605 | BD Biosciences |
| anti-mouse TER-119 | TER119 | FITC | Invitrogen |
| anti-mouse CD45 | 30-F11 | APC | BD Biosciences |
| anti-mouse CD71 | R17217 | PE | Invitrogen |
| Hoeschst |  |  | Sigma |
| anti-mouse Fc | 2.4G2 |  | BD Biosciences |
| anti-BrdU | 3D4 | FITC | Biolegend |

### Enzyme-linked immunosorbent assay (ELISA)

96 well plates (Corning) were coated with 2*μ*g of NANP9 or *Pf*16 peptide (Biomatik) diluted in PBS and incubated at 4°C overnight. Plates were washed with PBST (PBS with 0.05% Tween-20) and incubated for 1 hour at room temperature with blocking buffer (PBST + 3% milk). Plasma from murine samples were serially diluted in PBST+1% milk in duplicate, added to plates, and incubated at room temperature for 2 hours. Plates were incubated in anti-mouse IgG-HRP (Southern biotech) or anti-mouse IgM-HRP (Southern biotech) secondary antibody diluted at 1:2000 in PBST+1% milk for 1 hour. The plates were detected using 1X 3,30,5,50-Tetramethylbenzidine (TMB) (Invitrogen) and quenched with 1M HCl after 1.5 minutes. Optical density was measured using a spectrophotometer at 450nm and 570nm.

### Statistical analysis

Statistical analysis was performed by GraphPad Prism Software (La Jolla, CA) and analyzed via Mann Whitney *U* test, paired t test, or Wilcoxon matched-pair signed rank test as indicated.

## Acknowledgments

The authors would like to express sincere gratitude to all study participants in the IMRAS and BSPZV2 Research studies, which were essential to the success of this research. Flow cytometry data was acquired through the University of Washington Cell Analysis Facility Shared Resource Lab, with NIH award 1S10OD024979-01A1 funding for the Symphony A3. This research was supported by the National Institutes of Health grants F32AI178963 (C.E.M) and U01AI142001 (M.P) and Bill and Melinda Gates Foundation grants INV-010680, INV-043758, and INV-009411 (N.P.K).

---

**Extended Data Fig. 1:**
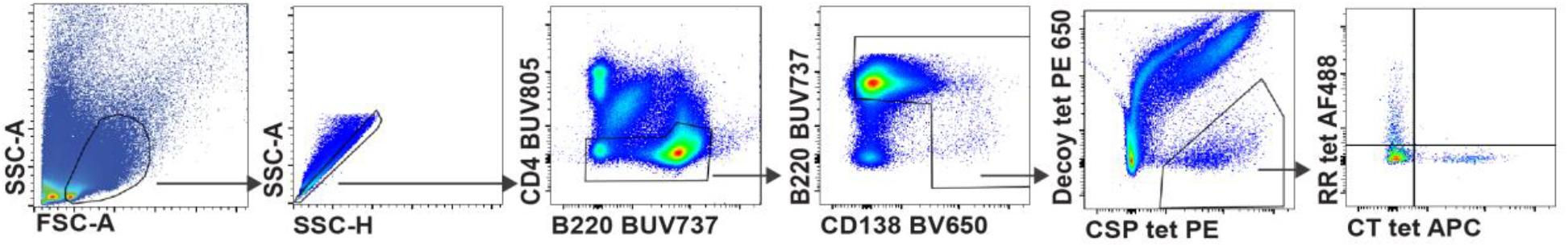
Representative gating on murine RR- and CT-specific B cells. Shown is the gating strategy to identify RR- and CT-specific B cells in a C57BL/6 mouse infected with 2000 *Py^Pf^*^CSP^ fourteen days prior. Gated on live lymphocytes, singlets, CD4^-^, B cells (B220^+^ or CD138^+^), full-length CSP^+^, RR^+^ or CT^+^.

**Extended Data Fig. 2:**
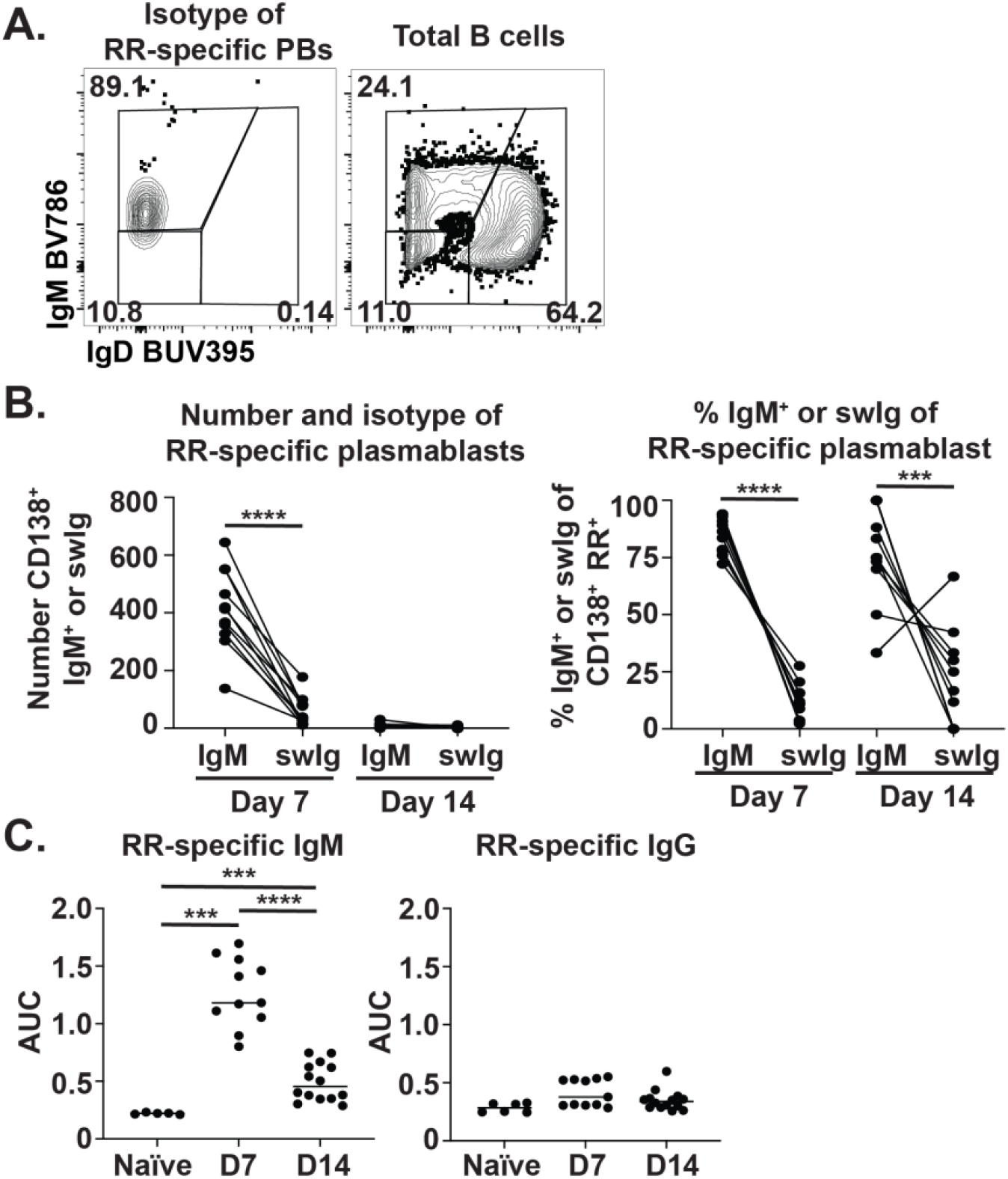
*Plasmodium* infection generates RR-specific IgM plasmablasts. C57BL/6 mice were infected with 2000 *Py^Pf^*^CSP^ sporozoites as in Fig. 1. Serum and spleens were harvested 7 or 14 days later. **a,** The isotype of the RR-specific PBs at days 7 post infection. Expression of IgD and IgM on total B cells is shown as a staining control. **b,** The number and proportion of RR-specific IgM or class-switched (swIg) PBs at indicated timepoints post infection. **c,** RR-specific serum IgM or IgG at indicated timepoints post infection. n=5-14 mice per time point. Statistical analysis by paired t test (b) or Mann-Whitney (c) ***p<0.005 ****p<0.0001

**Extended Data Fig. 3:**
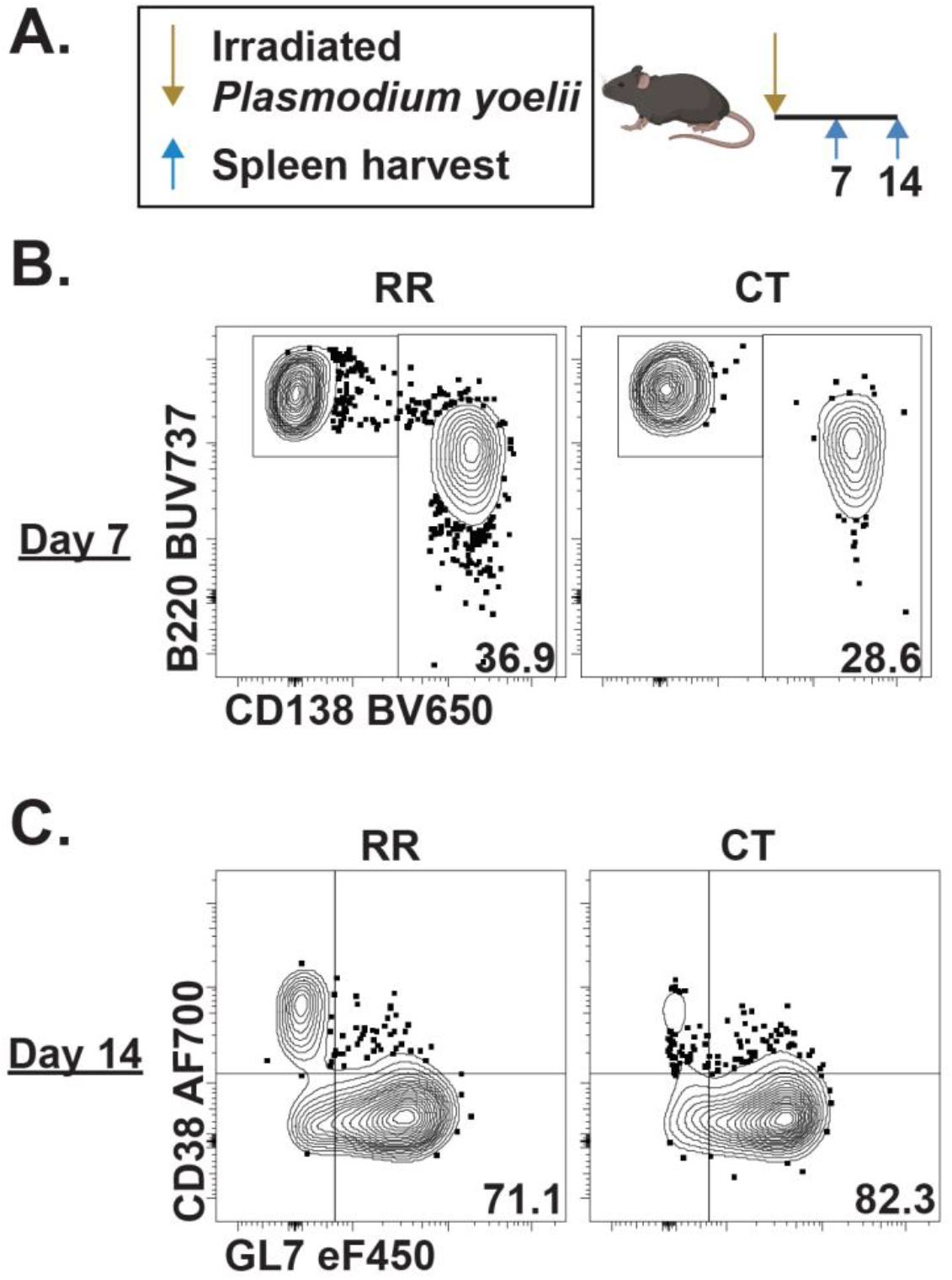
Immunization with irradiated *Plasmodium* generates RR-and CT-specific plasmablasts and germinal center B cells. **a,** C57BL/6 mice were immunized with 50000 irradiated *Py^Pf^*^CSP^ sporozoites intravenously. Spleens were harvested 7 or 14 days later. **b,** RR- and CT-specific PBs (CD138^+^) at day 7 post sporozoite immunization. **c,** RR- and CT-specific GCs (CD38^-^GL7^+^) 14 days post sporozoite immunization.

**Extended Fig. 4:**
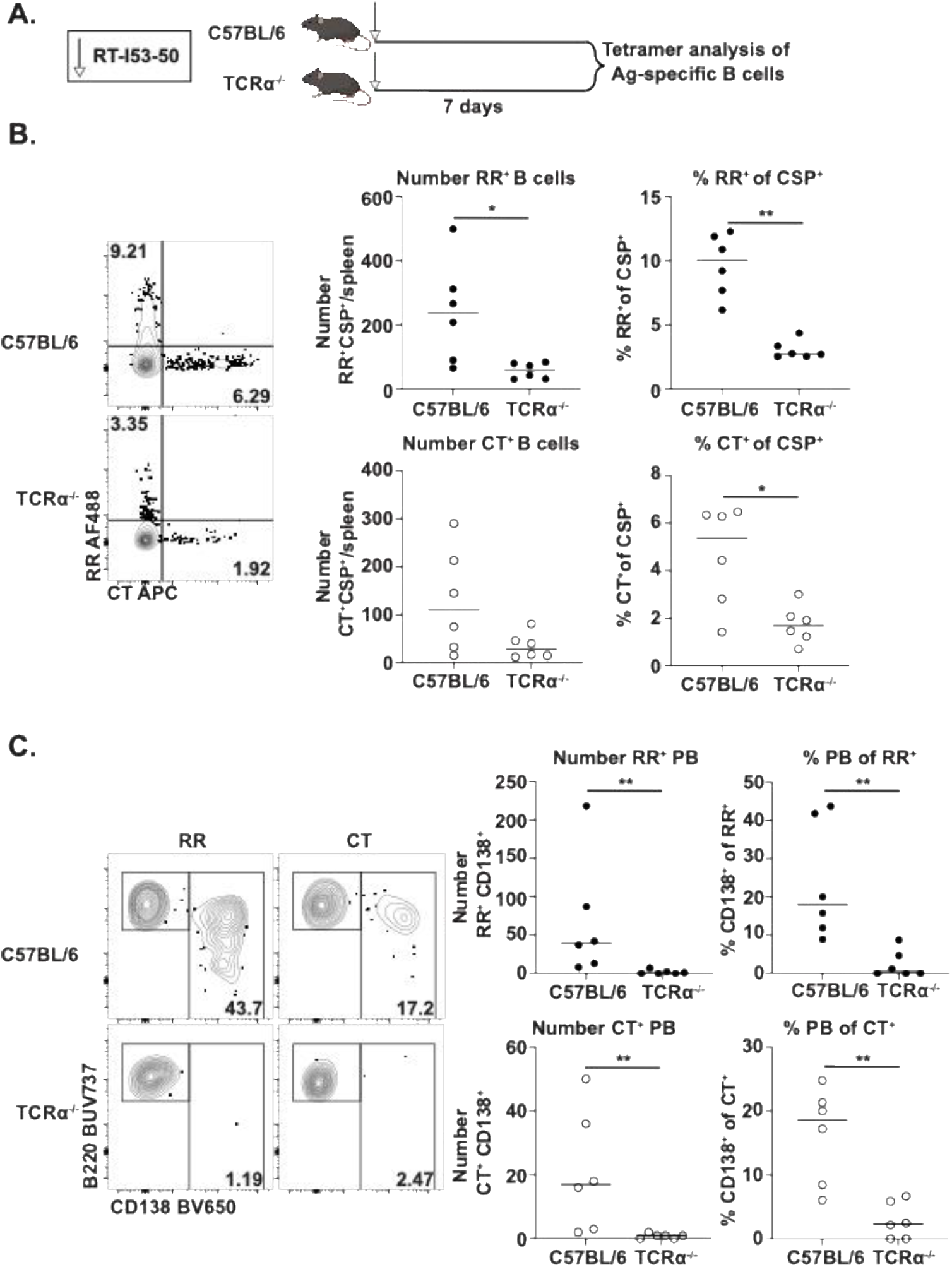
RR- and CT-specific responses are T-dependent. **a,** C57BL/6 or TCRα^-/-^ mice were immunized with 25*µ*g RT-I53-50 intraperitoneally. CT- and RR- specific B cells were isolated and characterized 7 days later. **b,** Number and proportion of RR- and CT-specific B cells. **c,** Number and proportion of RR- and CT-specific PBs (CD138^+^). n = 6. Statistical analysis by Mann–Whitney *p<0.05, **p<0.005

**Extended Data Fig. 5:**
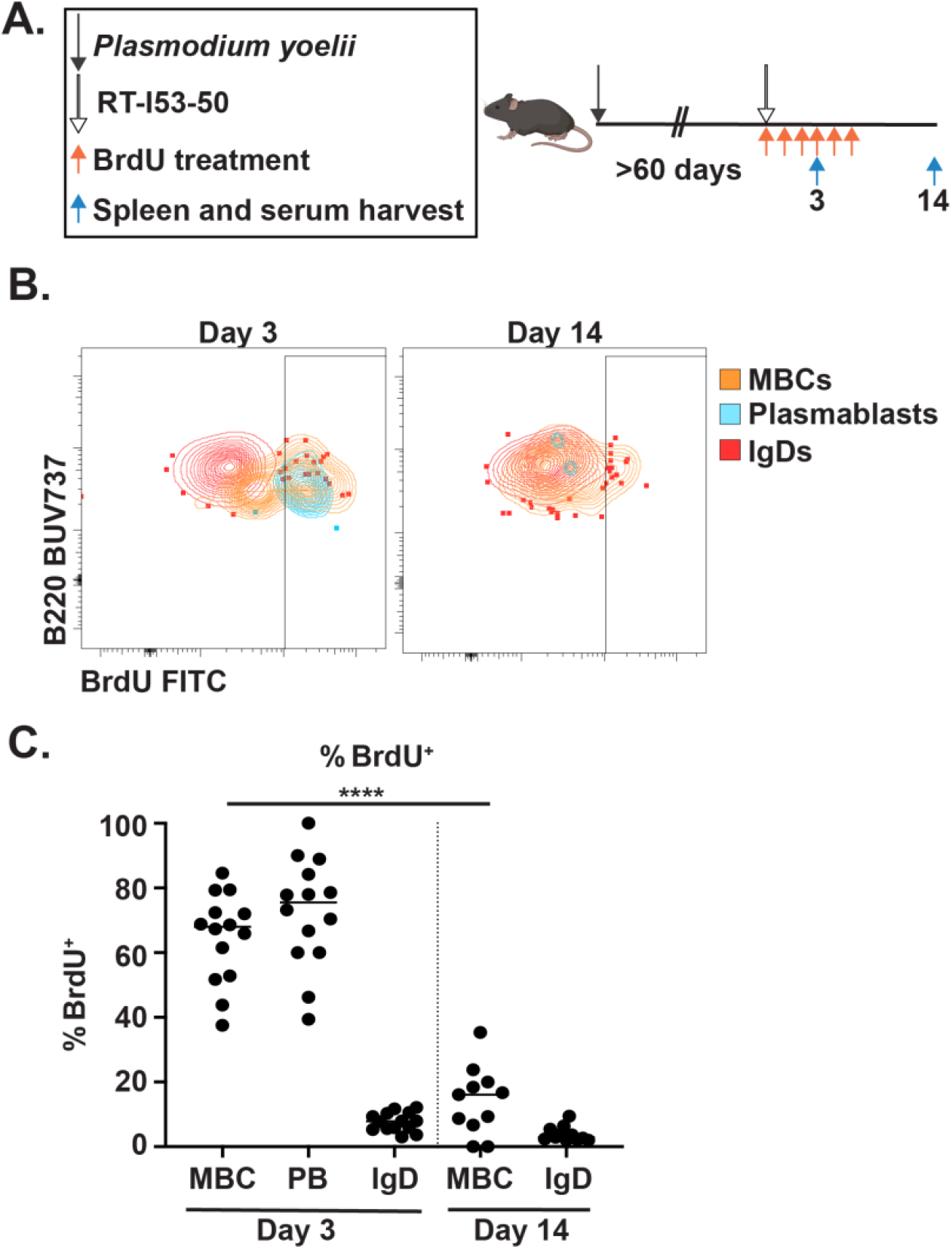
RR-specific MBCs do not proliferate into new, long-lived effectors. **a,** C57BL/6 mice were infected with 2000 *Py^Pf^*^CSP^. At least 60 days later, mice were immunized with 25*µ*g RT-I53-50 with concurrent BrdU treatment as indicated. Splenic RR-specific B cells were isolated and phenotyped. **b,** BrdU staining on RR-specific PBs (CD138^+^), MBCs (IgD^-^IgM^-^ CD73^+^CD80^+^), or IgD^+^ B cells. **c,** Proportion of RR-specific MBCs, PBs, or IgD^+^ B cells that are BrdU^+^. n = 6-14. Statistical analysis by Mann-Whitney ****p<0.0001

**Extended Data Fig. 6:**
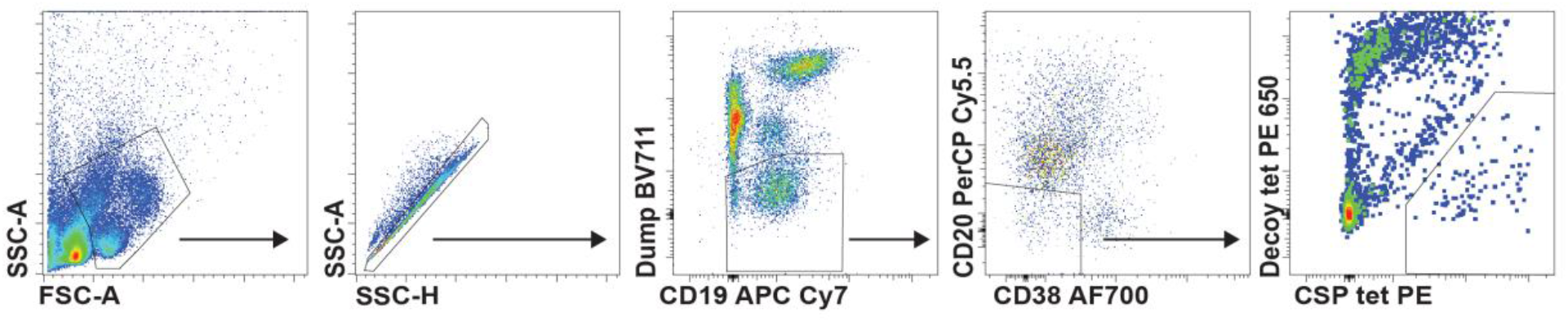
Representative gating for human samples. Shown is the gating strategy to identify antigen-specific cells in humans. Gated on live lymphocytes, singlets, dump^-^ (CD14, CD16, CD3), B cells (CD20^+^ or CD38^+^), full-length CSP^+^.

## Notes

### Competing Interest Statement

The authors have declared no competing interest.

### Summary of Updates

Reordered figures and made textual changes to enhance clarity

## References

1. Malaria vaccines (RTS,S and R21). *World Health Organization* https://www.who.int/news-room/questions-and-answers/item/q-a-on-rts-s-malaria-vaccine.

2. Wadman, M. First malaria vaccine slashes childhood deaths. Science (1979). 282, 357 (2023).

3. Olotu, A. et al. Seven-Year Efficacy of RTS,S/AS01 Malaria Vaccine among Young African Children. New England Journal of Medicine 374, 2519–2529 (2016).

4. RTSS Clinical Trials Partnership. Efficacy and safety of RTS,S/AS01 malaria vaccine with or without a booster dose in infants and children in Africa: final results of a phase 3, individually randomised, controlled trial. The Lancet 386, 31–45 (2015).

5. Agnandji, S. T. et al. Efficacy and Safety of the RTS,S/AS01 Malaria Vaccine during 18 Months after Vaccination: A Phase 3 Randomized, Controlled Trial in Children and Young Infants at 11 African Sites. PLoS Med. 11, e1001685 (2014).

6. Ali, M. S. et al. The anti-circumsporozoite antibody response to repeated, seasonal booster doses of the malaria vaccine RTS,S/AS01E. npj Vaccines *2025 10:1* 10, 1–11 (2025).

7. Gotuzzo, E., Yactayo, S. & Córdova, E. Efficacy and Duration of Immunity after Yellow Fever Vaccination: Systematic Review on the Need for a Booster Every 10 Years. Am. J. Trop. Med. Hyg. 89, 434 (2013).

8. Taub, D. D. et al. Immunity from Smallpox Vaccine Persists for Decades: A Longitudinal Study. Am. J. Med. 121, 1058 (2008).

9. Jongo, S. A. et al. Safety, Immunogenicity, and Protective Efficacy against Controlled Human Malaria Infection of Plasmodium falciparum Sporozoite Vaccine in Tanzanian Adults. Am. J. Trop. Med. Hyg. 99, 338–349 (2018).

10. Chatterjee, D. et al. Avid binding by B cells to the Plasmodium circumsporozoite protein repeat suppresses responses to protective subdominant epitopes. Cell Rep. 35, 108996 (2021).

11. Zavala, F., Cochrane, A. H., Nardin, E. H., Nussenzweig, R. S. & Nussenzweig, V. Circumsporozoite proteins of malaria parasites contain a single immunodominant region with two or more identical epitopes. Journal of Experimental Medicine 157, 1947–1957 (1983).

12. Zavala, F. et al. Rationale for Development of a Synthetic Vaccine Against Plasmodium falciparum Malaria. Science (1979). 228, 1436–1440 (1985).

13. Guttinger, M. et al. Human T cells recognize polymorphic and non-polymorphic regions of the Plasmodium falciparum circumsporozoite protein. EMBO J. 7, 2555–2558 (1988).

14. Kumar, S. et al. Cytotoxic T cells specific for the circumsporozoite protein of Plasmodium falciparum. Nature 1988 334:6179 334, 258–260 (1988).

15. Sinigaglia, F. et al. A malaria T-cell epitope recognized in association with most mouse and human MHC class II molecules. Nature 1988 336:6201 336, 778–780 (1988).

16. Dobaño, C. et al. Concentration and avidity of antibodies to different circumsporozoite epitopes correlate with RTS,S/AS01E malaria vaccine efficacy. Nat. Commun. 10, (2019).

17. Chaudhury, S. et al. Breadth of humoral immune responses to the C-terminus of the circumsporozoite protein is associated with protective efficacy induced by the RTS,S malaria vaccine. Vaccine 39, 968–975 (2021).

18. Tan, J. et al. A public antibody lineage that potently inhibits malaria infection through dual binding to the circumsporozoite protein. Nature Medicine 2018 24:4 24, 401–407 (2018).

19. White, M. T. et al. Immunogenicity of the RTS,S/AS01 malaria vaccine and implications for duration of vaccine efficacy: Secondary analysis of data from a phase 3 randomised controlled trial. Lancet Infect. Dis. 15, 1450–1458 (2015).

20. Schofield, L. The circumsporozoite protein of Plasmodium: a mechanism of immune evasion by the malaria parasite? WHO Bulletin OMS 68, (1990).

21. Fisher, C. R. et al. T-dependent B cell responses to Plasmodium induce antibodies that form a high-avidity multivalent complex with the circumsporozoite protein. PLoS Pathog. 13, e1006469 (2017).

22. Brooks, J. F. et al. Molecular basis for potent B cell responses to antigen displayed on particles of viral size. Nat. Immunol. 24, 1762 (2023).

23. Kato, Y. et al. Multifaceted Effects of Antigen Valency on B Cell Response Composition and Differentiation In Vivo. Immunity 53, 548 (2020).

24. Clutterbuck, E. A. et al. Pneumococcal Conjugate and Plain Polysaccharide Vaccines Have Divergent Effects on Antigen-Specific B Cells. J. Infect. Dis. 205, 1408 (2012).

25. Zhang, M. et al. A highly infectious Plasmodium yoelii parasite, bearing Plasmodium falciparum circumsporozoite protein. Malar. J. 15, (2016).

26. Keitany, G. J. et al. Blood Stage Malaria Disrupts Humoral Immunity to the Pre-erythrocytic Stage Circumsporozoite Protein. Cell Rep. 17, 3193–3205 (2016).

27. Zuccarino-Catania, G. V. et al. CD80 and PD-L2 define functionally distinct memory B cell subsets that are independent of antibody isotype. Nature Immunology 2014 15:7 15, 631–637 (2014).

28. Dogan, I. et al. Multiple layers of B cell memory with different effector functions. Nat. Immunol. 10, 1292–1299 (2009).

29. Pape, K. A., Taylor, J. J., Maul, R. W., Gearhart, P. J. & Jenkins, M. K. Different B cell populations mediate early and late memory during an endogenous immune response. Science 331, 1203 (2011).

30. Callahan, D. et al. Memory B cell subsets have divergent developmental origins that are coupled to distinct imprinted epigenetic states. Nat. Immunol. 25, 562–575 (2024).

31. Tomayko, M. M., Steinel, N. C., Anderson, S. M. & Shlomchik, M. J. Hierarchy of Maturity of Murine Memory B Cell Subsets. Journal of Immunology 185, 7146–7150 (2010).

32. Marcandalli, J. et al. Induction of Potent Neutralizing Antibody Responses by a Designed Protein Nanoparticle Vaccine for Respiratory Syncytial Virus. Cell 176, 1420 (2019).

33. Bale, J. B. et al. Accurate design of megadalton-scale two-component icosahedral protein complexes. Science 353, 389 (2016).

34. Langowski, M. D. et al. Elicitation of liver-stage immunity by nanoparticle immunogens displaying P. falciparum CSP-derived antigens. NPJ Vaccines 10, 1–14 (2025).

35. McNamara, H. A. et al. Splenic Dendritic Cells and Macrophages Drive B Cells to Adopt a Plasmablast Cell Fate. Front. Immunol. 13, 825207 (2022).

36. Mesin, L. et al. Restricted Clonality and Limited Germinal Center Reentry Characterize Memory B Cell Reactivation by Boosting. Cell 180, 92 (2020).

37. Tan, J. et al. A public antibody lineage that potently inhibits malaria infection through dual binding to the circumsporozoite protein. Nature Medicine 2018 24:4 24, 401–407 (2018).

38. White, M. T. et al. Immunogenicity of the RTS,S/AS01 malaria vaccine and implications for duration of vaccine efficacy: Secondary analysis of data from a phase 3 randomised controlled trial. Lancet Infect. Dis. 15, 1450–1458 (2015).

39. Lara-Escandell, M. et al. Duration and association with protection of NANP-repeat-specific and C-terminus-specific anti-circumsporozoite protein IgG responses following RTS,S/AS01E vaccination: an observational ancillary immunological study of a phase 3 clinical trial. Lancet Infect. Dis. 26, 832 (2026).

40. Scally, S. W. et al. Rare PfCSP C-terminal antibodies induced by live sporozoite vaccination are ineffective against malaria infection. Journal of Experimental Medicine 215, 63–75 (2018).

41. Wang, L. T. et al. Protective effects of combining monoclonal antibodies and vaccines against the Plasmodium falciparum circumsporozoite protein. PLoS Pathog. 17, e1010133 (2021).

42. Paus, D. et al. Antigen recognition strength regulates the choice between extrafollicular plasma cell and germinal center B cell differentiation. Journal of Experimental Medicine 203, 1081–1091 (2006).

43. Jongo, S. A. et al. Safety and Differential Antibody and T-Cell Responses to the Plasmodium falciparum Sporozoite Malaria Vaccine, PfSPZ Vaccine, by Age in Tanzanian Adults, Adolescents, Children, and Infants. Am. J. Trop. Med. Hyg. 100, 1433–1444 (2019).

44. McNamara, H. A. et al. Antibody Feedback Limits the Expansion of B Cell Responses to Malaria Vaccination but Drives Diversification of the Humoral Response. Cell Host Microbe 28, 572–585.e7 (2020).

45. Yamba, E. I. et al. Monthly Entomological Inoculation Rate Data for Studying the Seasonality of Malaria Transmission in Africa. MDPI 5, (2020).

46. Shaukat, A. M., Breman, J. G. & McKenzie, F. E. Using the entomological inoculation rate to assess the impact of vector control on malaria parasite transmission and elimination. Malar. J. 9, 122 (2010).

47. Hickey, B. et al. IMRAS—A clinical trial of mosquito-bite immunization with live, radiation-attenuated P. falciparum sporozoites: Impact of immunization parameters on protective efficacy and generation of a repository of immunologic reagents. PLoS One 15, e0233840 (2020).

48. Papadatou, I., Tzovara, I. & Licciardi, P. V. The Role of Serotype-Specific Immunological Memory in Pneumococcal Vaccination: Current Knowledge and Future Prospects. Vaccines (Basel). 7, 13 (2019).

49. Datoo, M. S. et al. Safety and efficacy of malaria vaccine candidate R21/Matrix-M in African children: a multicentre, double-blind, randomised, phase 3 trial. The Lancet 403, 533–544 (2024).

50. Tas, J. M. J. et al. Antibodies from primary humoral responses modulate the recruitment of naive B cells during secondary responses. Immunity 55, 1856–1871.e6 (2022).

51. Slifka, M. K. & Ahmed, R. Long-lived plasma cells: a mechanism for maintaining persistent antibody production. Curr. Opin. Immunol. 10, 252–258 (1998).

52. Taylor, J. J. et al. Deletion and anergy of polyclonal B cells specific for ubiquitous membrane-bound self-antigen. Journal of Experimental Medicine 209, 2065–2077 (2012).

53. Wargacki, A. J. et al. Complete and cooperative in vitro assembly of computationally designed self-assembling protein nanomaterials. Nature Communications 2021 12:1 12, 1–14 (2021).

